# Maternal smoking in early pregnancy disrupts placental function through syncytiotrophoblast and macrophage dysregulation

**DOI:** 10.1101/2025.07.10.664172

**Authors:** Daniela S. Valdes, Jose Nimo, Olivia Nonn, Juliane Ulrich, Désirée Forstner, Sandra Haider, Miriam Ressler, Marc Pignitter, Dominik N. Müller, Ralf Dechend, Martin Gauster, Fabian Coscia, Florian Herse

## Abstract

Smoking in pregnancy is the leading avoidable cause of gestational morbidity and mortality, causally linked to fetal growth restriction (FGR). The placenta, functional interface between mother and fetus is essential for healthy fetal development. For the first time, we studied cell type-resolved smoking effects on placental development at high molecular resolution using single-nucleus RNA sequencing and deep visual proteomics of matched tissues. We validated our findings through an independent cohort and in-vitro cigarette smoke exposure to primary human trophoblast cells. Our results show placental macrophages (Hofbauer cells; HBC) and the syncytiotrophoblast (STB) barrier are most affected by smoking, with dysregulation of cell-cell adhesion, extracellular matrix organization, and stress phenotype. STBs show moderate compositional increases in smokers and in-silico trophoblast differentiation modelling indicates a preferential shift towards the STB lineage in this group. The trophoblast displays a large upregulation of pro-angiogenic effectors, increases in xenobiotic detoxification, reduced mitochondrial function, and vastly altered transmembrane transport. These molecular changes affect placental development with important consequences for fetal growth. We provide insight into placental dysfunction contributing to FGR early in pregnancy, before clinical symptoms appear. We anticipate this data to advance diagnostics and therapies to improve FGR outcomes.

## Introduction

The placenta constitutes the interface between mother and fetus^1^. It regulates nutrient, metabolite and gas exchange, has an endocrine activity and thereby modulates maternal metabolism in pregnancy^2–4^. Healthy development and function of the placenta are essential for a successful pregnancy and fetal health, whilst developmental abnormalities modulate various disorders of pregnancy^5–7^.

Placental development occurs primarily in the first trimester of pregnancy. The most abundant phenotypes in the placenta are trophoblast cells. Mononucleated cytotrophoblasts (CTBs) form tree-like villi structures covered by a multinuclear syncytiotrophoblast (STB) layer^2,8^. The transcriptionally active STB establish the functional interface between maternal blood and placental villi. Due to its direct contact to maternal blood, the STB is involved in most functions of the placenta, as reflected by its high metabolic rate of up to 40% of the total feto-placental oxygen consumption^9^. Placental derived circulating factors are primarily secreted by the STB syncytium and have profound effects both on placental development and on maternal physiology^10,11^. Placental growth factor (PlGF), a member of the vascular endothelial growth factor (VEGF) family, is an important pro-angiogenic marker, and its imbalance is closely linked to obstetric disease^12^. In early pregnancy, it plays critical roles in placental development, in the context of vasculogenesis, angiogenesis and trophoblast lineage development^13,14^.

Adequate fetal development and growth are limited by the rate at which the placenta can deliver nutrients, oxygen and remove waste. Placental dysfunction has been associated with the pathophysiology like fetal growth restriction (FGR), pre-term labor, preeclampsia and miscarriage^15–17^. Importantly, detrimental effects extend well beyond pregnancy and affect the life-long health of both mother and offspring related to cardiovascular and metabolic disorders^6^. FGR is a leading cause of fetal morbidity and mortality worldwide^18^; which arises as a result of inadequate nutrient or oxygen supply at the maternal-fetal interface^19^.

Maternal smoking in pregnancy has a self-reported global prevalence of 1.7%, with a much higher European region estimate of 8.1%^20^. Of note, self-reporting underestimates true smoking status by up to 25% in Europe^21–23^. Smoking during pregnancy is associated with various pregnancy complications including placental abruption, placenta previa, ectopic pregnancy, preterm birth, FGR and low birthweight^24–26^. FGR, preterm birth and low birthweight are causally linked to maternal smoking^27–29^. Fetal birth defects in multiple organ systems are positively associated with maternal smoking^30^. At term, morphological effects of maternal smoking are well characterized and include a higher placental weight irrespective of smoking cessation in pregnancy^31^; increased calcifications and stiffened matrix^32,33^, reduced volume of intervillous space^34^ accompanied by profound adaptive angiogenesis in placental blood vessels within the villous mesenchymal core^35,36^. Additionally, functional effects include altered amino acid and glucose transport^37,38^, progesterone synthesis^39^ and estrogen metabolism^40^.

The placenta suffers oxidative stress and damage after sustained exposure to toxic smoke components in the circulation of maternal smokers. Reactive oxygen species (ROS) are associated with increased lipid peroxidation in the placenta later in pregnancy, with effects in cell membrane integrity and transport function^41^. Gene expression of *CYP1A1* and *CYP1B1* phase I xenobiotic metabolizing enzymes of the cytochrome P450 system is sharply upregulated in placentas from maternal smokers at term^42,43^, contributing to intracellular ROS formation. These are downstream of the activation of the aryl hydrocarbon receptor (AhR) pathway, well known for its activity in xenobiotic response^44^.

Maternal smoking in pregnancy is deleterious for placental health in every trimester of pregnancy, with reports largely focusing on global tissue dysregulation in the third trimester with limited cell phenotype specificity. However, we currently lack a detailed understanding of the transcriptional and proteomic changes of maternal smoking in first trimester placental tissue. Herein, we profiled the transcriptomic and proteomic landscape of the early human placenta to characterize the profound impact of smoking at the single-phenotype level during the developing phase of the placenta early in pregnancy. We describe molecular origins of aberrant interface capacity that likely mechanistically underpin FGR and other adverse obstetric outcomes later in pregnancy.

## Results

### Maternal smoking effects in early pregnancy are driven by placental barrier and immune capacity dysregulation

To comprehensively understand phenotype-specific molecular effects of maternal smoking on the developing human placenta, we analyzed first trimester tissues from elective terminations (5-11 weeks of gestation) (**Fig 1A**). We applied 10X Genomics single-nucleus RNA sequencing (snRNAseq) to high-quality nuclei isolated from 11 placentas (*n* = 5 smokers, *n* = 6 non-smokers); and conducted spatially resolved deep visual proteomics (DVP)^45^ of formalin-fixed paraffin-embedded (FFPE) villi from a subset of the same tissues (*n* = 3 smokers, *n* = 3 non-smokers). Importantly, these methods allowed for the unique and extensive molecular profiling of the STB, seldom performed since this syncytium is impossible to derive into a single cell suspension. Samples were matched for gestational age, maternal age and maternal body mass index (BMI) (**Suppl Fig 1, Methods Table 1**). In addition, we validated findings in an independent cohort (*n* = 23 smokers, *n* = 15 non-smokers) and through *in-vitro* functional assays using primary derived human trophoblast stem cells.

**Figure 1.**
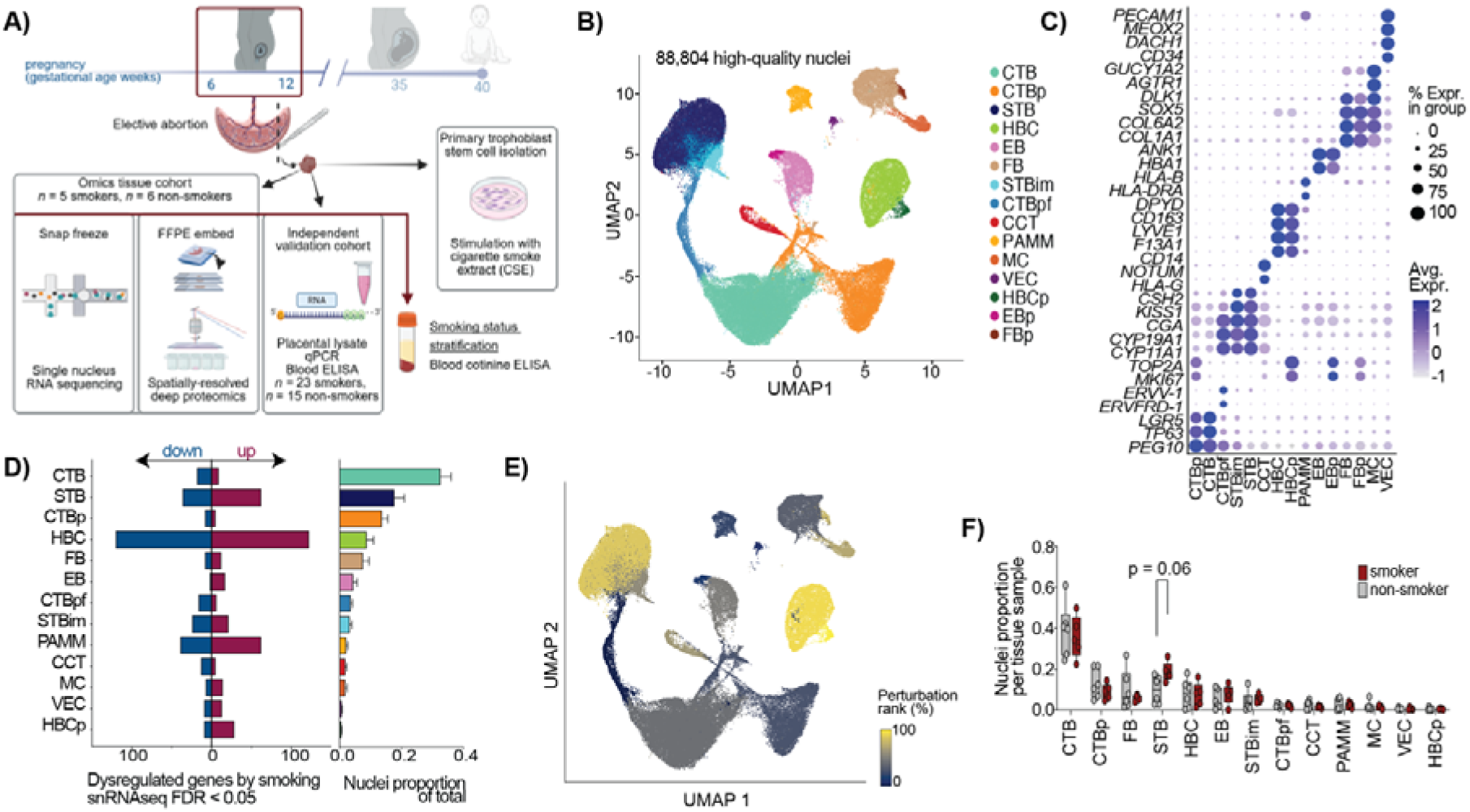
Maternal smoking induces phenotype specific gene expression dysregulation patterns, with the syncytiotrophoblast and Hofbauer cells being most affected. **(A)** Schematic representation of study design and samples used. Two main cohorts were included, the ‘omics cohort’ and an ‘independent validation cohort’, both of which stratified maternal smoking status based on serum cotinine levels and self-reporting. In addition, primary trophoblast cells isolated from self-reported non-smokers were used for functional in-vitro validations. FFPE, formalin-fixed paraffin-embedded; ELISA, enzyme linked immunosorbent assay; qPCR, quantitative polymerase chain reaction. Created with BioRender.com. **(B)** Uniform manifold approximation and projection (UMAP) visualization of harmonized dataset colored by the fifteen resolved and annotated cell types and states (*n* = 88,808 nuclei). CTB, cytotrophoblast; STB syncytiotrophoblast; CCT, cell column trophoblast; HBC, Hofbauer cell; PAMM, placental-associated maternal macrophage; EB, erythroblast; FB, fibroblast; MC, myocyte; VEC, vascular endothelial cell; p, proliferating; pf, pre-fusion; im, immature. **(C)** Dotplot visualizing mean gene expression of the canonical literature-based markers in the annotated groups. **(D)** Left: Number of differentially expressed genes by maternal smoking at a 10% FDR cut off (Upregulated genes (purple); downregulated genes (blue)). Dysregulation profiles are plotted in descending order based on the cell type or state’s contribution to the total tissue nuclei. Right: Bar plot of nuclei proportions of total per annotated cell type and state as an average of all included tissues, irrespective of maternal smoking status. Mean plotted with error bars representing the standard error of the mean. **(E)** Uniform manifold approximation and projection (UMAP) embedding of harmonized dataset colored by perturbation between maternal smoking status calculated using a machine learning classifier model. Color indicates perturbation rank percentage (%) based on area under the curve measurements, where higher values (yellow) indicate higher transcriptomic perturbation. **(F)** Box and whiskers plots of nuclei tissue proportions between smokers (red) and non-smokers (grey) per annotated cell type or state group. Each dot represents a placental tissue 25^th^ to 75^th^ replicate. Box extends from percentiles with a line at the median, whiskers extend from the minimum to maximum distribution values. Differences assessed by two-tailed unpaired Welch’s t-test.

**Table 1.** Characteristics of included study subjects.

|  | Omics cohort |  |  | Independent validation cohort<br>mRNA<br>PIGF ELISA |  |  |
| --- | --- | --- | --- | --- | --- | --- |
| Parameter | Non-smokers | Smokers | p | Non-smokers | Smokers | p |
| Sample size (n) | 11 | 5 | - | 14<br>24 | 24<br>36 | -<br>- |
| Cotinine (ng/mL) | n.d. | 147.2 ± 42.1 | 0.002* | n.d.<br>n.d. | 147.4 ± 153.5<br>105.2 ± 97.0 | <0.0001*<br><0.0001§ |
| Gestational age (days) | 57.8 ± 16.4 | 60.0 ± 9.4 | 0.8* | 61.8 ± 17.0<br>68.7 ± 15.5 | 59.6 ± 15.4<br>6.0 ± 14.5 | 0.6*<br>0.5* |
| Placental volume (cm<sup>2</sup>) | 1.6 ± 1.6 | 2.2 ± 1.2 | 0.5* | 3.2 ± 2.4<br>5.4 ± 8.9 | 2.6 ± 1.8<br>4.9 ± 6.9 | 0.4*<br>0.7§ |
| Maternal age (years) | 28.0 ± 5.3 | 25.4 ± 4.2 | 0.4* | 34.7 ± 6.9<br>32.1 ± 7.7 | 26.3 ± 6.0<br>28.7 ± 6.9 | 0.002*<br>0.09* |
| Maternal BMI (kg/m<sup>2</sup>) | 21.2 ± 2.7 | 22.4 ± 2.6 | 0.5* | 25.3 ± 3.7<br>25.2 ± 5.4 | 21.9 ± 2.8<br>22.3 ± 3.6 | 0.03*<br>0.03§ |
| Maternal weight (kg) | 60.0 ± 10.8 | 63.8 ± 4.1 | 0.5* | 69.8 ± 10.4<br>70.3 ± 15.9 | 60.8 ± 9.2<br>61.7 ± 12.2 | 0.01*<br>0.02§ |
| Maternal height (m) | 1.7 ± 0.05 | 1.7 ± 0.07 | 0.7* | 1.7 ± 0.09<br>1.7 ± 0.08 | 1.7 ± 0.06<br>1.7 ± 0.07 | 0.9*<br>0.7* |

Data presented as mean ± standard deviation. n.d. not detected, ELISA detection level was 1 ng/mL. Significance between groups evaluated by two-tailed unpaired Welch's t-test (\*) or two-tailed unpaired Mann Whitney test (§) depending on dataset normality and indicated with a symbol after the p-value. Additional samples were included in the validation cohort for PIGF ELISA measurement and characteristics noted. BMI, body mass index; PIGF, placental growth factor; ELISA, enzyme-linked immunosorbent assay.

Maternal smoking status was stratified by agreement between self-reporting and circulating cotinine levels, a stable metabolite from nicotine and clinical gold standard for assessing smoking status^46^. In the cohort used for snRNAseq and DVP techniques, cotinine cut-offs of > 50 ng/mL (heavy smokers, range 88–183 ng/mL) and < 1 ng/mL (non-smokers) were used to stratify samples, representing effects of heavy smoking on organ development and function. For the independent validation cohort, cotinine cut-offs of > 9 ng/mL (smokers, range 9-536 ng/mL) and <1 ng/mL (non-smokers) were used (**Suppl Fig 1**). After ambient RNA correction, doublet detection and stringent quality-control filtering, we retained 88,808 high-quality nuclei (*n* = 38,756 smokers, *n* = 50,052 non-smokers) for subsequent analyses (**Methods**). Sample normalization, batch correction and integration methods were applied to gain a harmonized dataset unconfounded by technical or biological sources of variance (**Suppl Fig 1**).

Nine highly specific cellular phenotypes encompassing three functional cellular states were resolved and annotated using a combination of canonical markers specificity, differential and conserved gene expression between communities (**Fig 1B,C, Methods**). Six describe the trophoblast lineage, including the cytotrophoblast (CTB) and its proliferating (CTBp) and pre-fusion (CTBpf) states; the syncytiotrophoblast (STB) and its recently described immature state (STBim)^19^ and the cell column trophoblast (CCT). Three described the placental immune repertoire, including placental (Hofbauer cells, HBC) and its proliferating state (HBCp), as well as placental-associated maternal macrophages (PAMM)^47^. Mesenchymal populations included fibroblasts (FB) and their proliferating state (FBp) as well as placental myocytes (MC). Finally, vascular endothelial cells (VEC), erythroblasts (EB) and their proliferating state (EBp) were identified. Cell cycle phase analysis showed clear and distinct separation between annotated non-proliferating and proliferating populations, with the latter mapping to the G2M and S phase related genes (**Suppl Fig 2**). The most correlated transcriptional profiles were found amongst cell states of the same phenotype (**Suppl Fig 2**).

We evaluated the transcriptomic dysregulation of populations (> 500 nuclei) using a pseudo-bulk approach (edgeR) via a likelihood ratio test generalized linear model framework that accounted for fetal sex and sequencing batch as covariates and corrected for multiple comparisons (**Fig 1D left**). Fetal sex was inferred using sex-associated genes (**Suppl Fig 1**). With a 5% false discovery rate (FDR) threshold, macrophages in the dataset showed the largest magnitude of dysregulation, led by the HBC (7.6% of nuclei, 242 DEGs [122 up, 120 down]), PAMM (2.0% of nuclei, 101 DEG [62 up, 39 down]) and HBCp, (0.5% of nuclei, 37 DEGs [28 up, 9 down]) (**Fig 1D**). Within non-immune groups, the STB (19.1% of nuclei, 98 DEGs [62 up, 36 down]) and its immature state (STBim, 3.6% of nuclei, 45 DEGs [21 up, 24 down]) exhibited the largest dysregulation, despite the much higher tissue abundance of CTB (32.8% of nuclei, 27 DEGs [9 up, 18 down]) and its proliferating (CTBp, 14.6% of nuclei, 13 DEGs [5 up, 8 down]) and pre-fusion (CTBpf, 4.0% of nuclei, 22 DEGs [6 up, 16 down]) states. Transcriptomic dysregulation in other cell types and states include: VEC (0.7% of nuclei, 22 DEGs [13 up, 9 down]); MC (1.6% of nuclei, 21 DEGs [14 up, 7 down]); FB (6.6% of nuclei, 20 DEGs [12 up, 8 down]); EB (4.0% of nuclei, 19 DEGs [17 up, 2 down]) and CCT (2.1% of nuclei, 18 DEGs [5 up, 13 down]) (**Fig 1D, Suppl Table 2**). The high-magnitude dysregulation of the STB and HBC groups was validated using a machine-learning classifier framework that prioritizes and ranks cell types based on predicted biological perturbations in a high-dimensional space^48^ (**Fig 1E**). A higher percentage rank value (calculated based on predicted areas under the curve) indicates an increased biological perturbation. Interestingly, dysregulation proved to be phenotype specific (i.e. within a cell type and it is associated state(s)), without evident distinct global patterns (**Suppl Fig 2**). ADP ribosylation factor like GTPase 17A (*ARL17A*), associated with protein trafficking and transport, was the only protein-coding gene concomitantly upregulated in trophoblast (STB, STBim, CTBpf), immune (HBC) and mesenchymal (VEC, FB) groups (**Suppl Fig 2**). We found a moderate increase in STB nuclei in placentas from smokers (mean difference = 9.1% STB nuclei, *p* = 0.06) and no differences for any other cell type or state (**Fig 1F**). There was no evident depletion of a specific region of the STB cluster between smoking groups, indicating no enrichment of a specific transcriptomic profile for the higher STB nuclei numbers in smokers (**Suppl Fig 2**).

Together, we found widespread gene expression changes induced by maternal smoking in early pregnancy and identified the STB syncytium and HBC resident macrophages as most affected. These changes influence the development and function of this organ and underpin the pathophysiology of causally-related pregnancy complications to smoking.

To shed light into the regulatory mechanisms governing trophoblast and immune response to maternal smoking in early pregnancy as a proxy for placental health, we further investigated differential expression patterns of the STB and HBC, the most dysregulated cell types (**Fig 2A, D**). Interestingly, 9.5% of dysregulated STB genes map to Human Proteomic Atlas (HPA) annotated secreted markers, whilst 22.2% map to HPA annotated membrane markers (**Suppl Fig 3**). Similarly for the HBC, 8.3% of dysregulated genes map to secreted proteins, whilst 19% map to membrane proteins. The high proportion of dysregulated membrane and secreted genes point to compromised intercellular placental barrier communication and fetal-maternal exchange.

**Figure 2.**
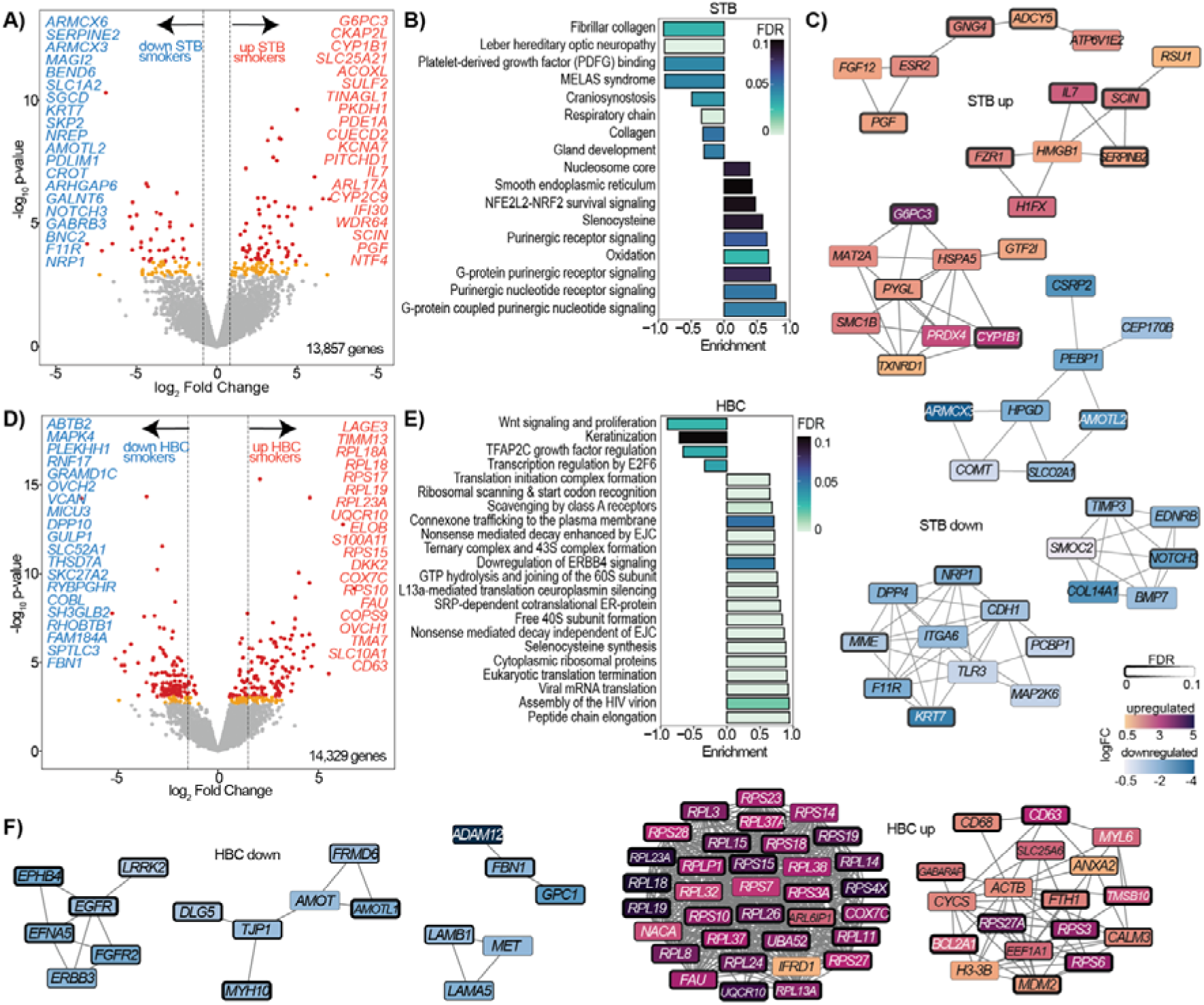
Syncytiotrophoblast and Hofbauer cell transcriptomic dysregulation and associated functionally enriched pathways and master modulators. **A, D)** Volcano plot of differential expressed genes (DEGs) analysed by snRNAseq from syncytiotrophoblast (STB) **(A)** and Hofbauer Hofbauer cells (HBC) **(D).** The top 20 most dysregulated genes are shown. DEGs were calculated using semi-bulk counts per cell type via a generalizer linear model and Benjamini-Hochberg correction in the edgeR package. **B, E)** Pathway enrichment analysis of dysregulated STB **(B)** and HBC **(E)** genes (absolute FC > 0.25, FDR < 0.1). Enrichment was done based on ranked log fold changes spanning the gene set enrichment analysis (GSEA), reactome and the WikiPathways databases. Positive enrichment values represent pathways enriched in smokers; negative values represent those enriched in non-smokers. Bar color is mapped to dysregulaiton FDR. **C, F)** Hub genes of upregulated (log2FC > 0.25, FDR < 0.1) and downregulated (log2FC < - 0.25, FDR < 0.1) tissue STB **(C)** and HBC **(F)** genes by maternal smoking in early pregnancy, based on protein-protein interaction background stringDB network followed by topological analysis.Color represents log2FC, node border thickness corresponds to differential expression analysis’ FDR.

The AhR xenobiotic detoxification response characterized by *CYP1B1* in the placenta was restricted to the STB phenotype, highlighting the importance of cell type resolved analyses (**Fig 2A**). Dysregulated STB genes suggested an impairment in placental barrier integrity. In addition to dysregulation of important membrane components (*PHLDB2, ITGA6, SGCD, DSC3, SDK1, F11R*) there was a myriad of dysregulated transporters for glutamate (*SLC1A2*), ions (*KCNA7, LRRC8B*), amino acids (*SLC1A4*), fatty acids (*SLC27A2*), sugars (*SLC45A4*), hormones (*SLC16A2, ESR2*), low-density lipoproteins (*LRP2*), prostaglandins (*SLCO2A1*) nucleosides (*SLC28A1*) and nucleotide sugars (*SLC35B4*) (**Suppl Table 2**). In addition, there was a marked upregulation of pro-angiogenic placental growth factor (*PGF*) and pro-vasculogenic adrenomedullin (*ADM*) that play critical roles in placental vascular and trophoblast development. Functional pathway analysis described impaired mitochondrial function and tissue remodeling of the STB, with a concomitant induction in oxidative stress and extracellular signaling response in smokers. To better understand the complex interactions between dysregulated genes we performed network analysis and identified hub genes, defined as those with the highest connectivity across DEGs, which represent genes that play central roles in the network, as either potential regulators or bottlenecks in biological pathways (Figures 2C**, F, Methods**). Identified markers clustered into functionally coherent communities (**Fig 2C**). Upregulated STB hub genes formed three distinct communities, associated with metabolic regulation and redox response; immune regulation and cytoskeletal structure; as well as angiogenesis and signal transduction. Downregulated STB hub genes also formed three distinct communities, associated with cell adhesion and metabolism; regulation of cytoskeletal organization and cell proliferation; and ECM remodeling and cell signaling. We predicted transcription factors (TFs) modulating hub gene dysregulation networks and found major players of trophoblast development (*EP300, GATA3, TEAD4, JUND*) (**Suppl Fig 3**).

In the HBC, transcriptomic dysregulation is consistent with hallmarks of macrophage activation and a pro-inflammatory state, highlighted by increases in phagocytic (*CD14, CD63, CD68*), and stress response (*S100A11, BCL2A1, MDM2*) markers (**Fig 2D**). Functional pathway enrichment showed reduced signaling in Wnt/ERBB4 pathways and TFAP2C growth factor regulation, important players in physiological placental morphogenesis and development (**Fig 2E**). In addition, a multitude of pathways associated with ribosomal biogenesis and translational capacity were upregulated in HBCs of smokers, consistent with heightened biosynthetic demands of activated macrophages. Upregulated hub gene communities in the HBC delineate main modulators of this pro-inflammatory and stress-adapted transcriptional state associated with metabolic activation and increased protein synthesis. Downregulated hub genes clustered into communities representative of tight junction and cytoskeletal regulation, ECM remodeling and growth factor signaling. In line with these findings, TFs predicted to coordinate this dysregulation are linked to basal and stress-inducible transcriptional regulation (*TAF1, TAF7, POLR2A, TP53*) (**Suppl Fig 3**).

These results demonstrate a striking cell type specific transcriptional reprogramming of the STB and HBC in response to maternal smoking. Both cell types feature coordinated alterations in metabolic and structural programs capable of impairing organ development and function, with implications for fetal growth.

Trophoblast cell types differentiate in the first trimester of pregnancy to enact specialized functions that allow for nutrient and gas exchange between mother and fetus. We used primary derived trophoblast stem cells (TSC)^49–51^ to model human trophoblast function and investigated molecular effects of cigarette smoke components on the STB phenotype after *in-vitro* directed TSC fusion (**Fig 3**). For the first time, we additionally characterized the proteomic landscape of this model by high-sensitivity mass spectrometry (*n* = 8,837 protein IDs) (**Suppl Fig 4**), where we observed clear proteomic shifts between undifferentiated and differentiated states, encompassing known and novel phenotypic and functional protein markers.

**Figure 3.**
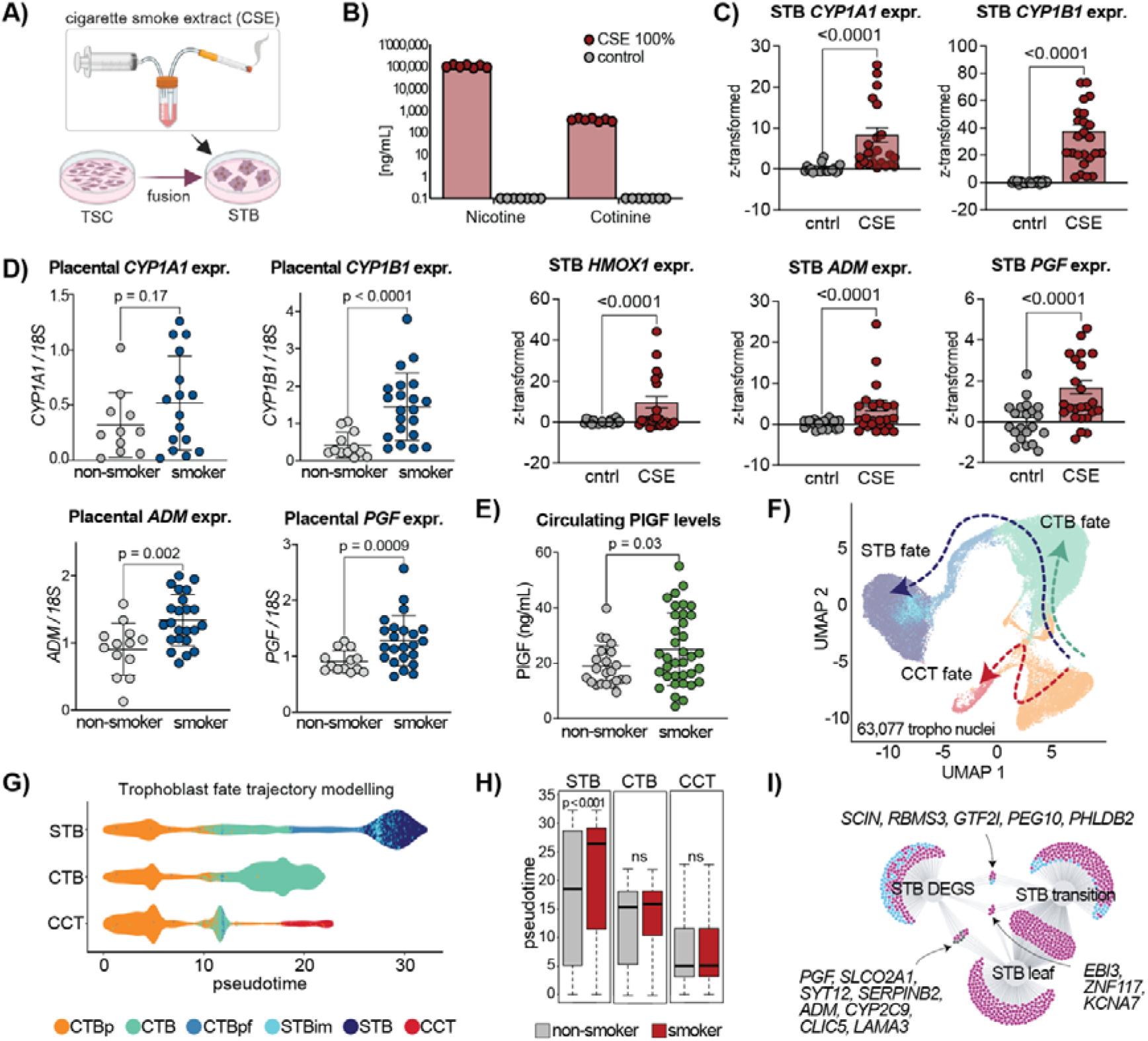
Validation of smoking effects on syncytiotrophoblasts xenobiotic stress and pro-angiogenic signature. **(A)** Schematic representation of cigarette smoke extract (CSE) preparation used for *in-vitro* stimulations as reported in Gellner et al^136^. Reference research cigarettes are bubbled through cell culture medium at a fixed rate. Resulting medium is acidic, so pH is adjusted and sterile filtered. For reproducibility, 100% CSE generated from 10 experiments were pooled into batches (n = 4 total batches). Created with biorender.com. **(B)** Cigarette smoke extract (CSE) media was validated by nicotine and cotinine levels analyzed by mass spectrometry. **(C)** Gene expression measured by qPCR in STB fused primary cells after 6h of CSE stimulation (n = 3 independent experiments with technical sextuplicates each). Data was normalized with a z-transformation per experiment as qPCR was measured in different plates. Significance assessed with unpaired two-tailed Mann-Whitney-tests. **(D)** Gene expression measured via quantitative PCR of whole placenta lysate of an independent cotinine-validated maternal smoking in early pregnancy cohort (*n* = 23 smokers, *n* = 15 non-smokers). Differences between groups assessed using two-tailed unpaired Welch t-tests. **(E)** Serum levels of placental growth factor in maternal serum measured via enzyme-linked immunosorbent assay (ELISA) in an independent cotinine-validated pregnancy cohort (*n* = 36 smokers, *n* = 22 non-smokers). Differences between groups were assessed using a two-tailed unpaired Welch t-test. **(F)** Uniform manifold approximation and projection (UMAP) embedding of trophoblast types and states used for trajectory modelling (*n* = 63,077 total nuclei; *n* = 27,055 nuclei from smokers, *n* = 36,022 nuclei from non-smokers). **(G)** Scatter plot of modelled pseudotime values separated by inferred lineage. Each dot represents a nucleus, colored by its annotated cell type or state membership. **(H)** Box plots of pseudotime distributions per lineage inferred (spanning from progenitor to endpoint) separated by maternal smoking status. Boxes represent interquartile range (IQR) from first to third quartile; median is indicated by the line inside the box. Whiskers extend to 1.5 times the IQR from each quartile. Significance between smokers and non-smokers for all three lineages was tested through a permutation, Kolmogorow-Smirnov, and generalized linear model tests. **(I)** Overlap between dysregulated genes in the STB phenotype and trajectory modelled STB leaf and transition genes. For the overlap, STB DEGs included those with an FDR < 0.1 and log2FC ± 0.25; transition genes included those with an FDR < 0.05 and absolute Spearman correlation of >0.4; leaf genes included those with an FDR < 0.05, logFC > 3 and log counts per million of > 4. Colours indicate agreement between upregulated (magenta) and downregulated (blue) markers. TSC, trophoblast stem cell; STB, syncytiotrophoblast; CTB, cytotrophoblast; CCT, cell column trophoblast; p, proliferating; pf, pre-fusion; im, immature.

Because tobacco smoke has over 7,000 constituents, cigarette smoke extract (CSE) containing nicotine, cotinine and other soluble constituents acts as a better model than nicotine alone to study the biological effects of smoking^52^. We generated standardized CSE using 1R6F reference small batch research cigarettes^53^ as previously described in the literature^54^, where cigarette smoke is bubbled through cell culture medium at a controlled rate (**Fig 3A**). We confirmed expected nicotine and cotinine concentrations in the CSE by mass spectrometry (**Fig 3B**). Additionally, we characterized volatile and semi-volatile constituents by GC-MS, as these act as potential ligands for smoke-related cellular responses (**Suppl Fig 5**).

At first, we validated the dysregulation of specific marker genes of STB physiology previously described by snRNAseq. After validating the time course (Suppl Fig 5), we stimulated STBs derived from TSCs with CSE for 6 hours. By qPCR, we revealed the induced expression of xenobiotic enzymes *CYP1A1* and *CYP1B1*, heme oxygenase-1 (*HMOX1*) *and important* pro-angiogenic *PGF* and *ADM* (**Fig 3C**). We further confirmed these findings in our independent validation cohort. Despite the STB only consisting of 19.1% of total placental nuclei, the upregulation of *CYP1B1*, *PGF* and *ADM* in the smoking group is still captured at the whole placental tissue lysate (**Fig 3D**). Importantly, we found the induction of *PGF* expression to have systemic implications, as circulating PlGF protein levels are elevated in the plasma of smoking mothers (mean difference 6.2 ± 2.7 ng/mL, *p* = 0.03) (**Fig 3E**).

In light of our findings of elevated STB nuclei proportions in smokers (**Fig 1F**) and the upregulation of pro-angiogenic PGF and ADM, with known roles in trophoblast development, we further investigated whether maternal smoking may impact healthy trophoblast development using pseudotime analysis *in-silico*. Here, cell differentiation dynamics are reconstructed based on individual nuclei gene expression fingerprints in combination with the high-dimensional UMAP space to identify global lineage structures (**Fig 3F-H**). Transcriptomic data from the nuclei of the six trophoblast cell types and states (CTBp, CTB, CTBpf, STB, STBim, CCT) was used for trajectory inference (*n* = 63,077 nuclei; *n* = 27,055 smokers, *n* = 36,022 non-smokers) (**Fig 3F, Suppl Fig 6**). To avoid cell annotation label bias, data was re-clustered (resolution 0.3 yielding 10 clusters), and clusters used for trajectory modelling (**Suppl Fig 6**). As a result, three biologically relevant lineages were inferred, starting from one of the two proliferating CTB (CTBp) subclusters, towards the multinucleated STB (STB-fate), towards the invasive CCT (CCT-fate) and between the proliferating and non-proliferating CTB (CTB-fate) (**Fig 3F, G, Suppl Fig 6, Suppl Table 3**).

To better understand trophoblast differentiation, key regulatory intermediates (transition) and terminal effector (leaf) genes were inferred using the non-smokers nuclei exclusively (*n* = 36,022 nuclei) (**Fig 3C, Methods**). Transition genes represent temporally expressed genes that change dynamically across pseudotime and are likely modulators of the differentiation process, whereas leaf genes demarcate the bifurcation and differentiated states.

When comparing trajectories between smokers and non-smokers, we found highly differential pseudotime distributions exclusively in the STB lineage (**Fig 3H**). This suggests a preferential differentiation towards the modelled STB lineage with a small effect size (odds ratio 1.02; i.e. 2% preference to the STB lineage per unit of pseudotime) in smokers compared to non-smokers. Albeit small, this increase appears to be functionally relevant, as transition and leaf genes closely linked to placental health are dysregulated in the STB (**Fig 3I**). Three genes dysregulated by smoking were identified as both leaf and transition markers, including *KCNA7, ZNF117* and *EBI3*, involved in ion transport, transcriptional regulation and immune modulation, respectively. Dysregulated transition genes were associated to cytoskeletal remodeling, trophoblast lineage specification and gene expression control, highlighting their putative role towards a skewed differentiation to the STB in smokers. Leaf genes overlapping with STB DEGs reflected disruptions in angiogenesis, cell adhesion, prostaglandin transport and ECM organization, all of importance for barrier function homeostasis.

We validated primordial snRNAseq findings of STB xenobiotic and pro-angiogenic responses on an independent whole placental lysate cohort and using *in-vitro* primary derived human trophoblast cells stimulated with soluble components of cigarette smoke through CSE medium prepared according to^54^. Trophoblast differentiation modeling revealed a subtle yet preferential shift towards the STB lineage in smokers. Smoking-associated dysregulation was linked to regulatory and effector genes governing cell signaling and cytoskeletal remodeling. These data support the notion that smoking disrupts core molecular programs of placental function and may influence trophoblast fate decisions.

### Deep visual proteomics reveals mitochondrial syncytiotrophoblast accumulation and hemostatic involvement

As next step to comprehensively understand the early pregnancy molecular dysregulation by maternal smoking, we applied deep visual proteomics (DVP)^45^ and collected single cells via laser microdissection followed by ultra-high sensitivity label-free mass-spectrometry based proteomics (**Fig 4A**). The main cell types of interest based on snRNAseq findings **(Fig 4B, C)** included HBCs that constitutively express cluster of differentiation CD163 (empty arrow), CTBs characterized by epithelial cadherin (E-cadherin, coded by the *CDH1*; filled arrow) and STBs identified by being E-cadherin negative and towards the outside of the villous structures (i.e. where maternal blood flows around the villi; dotted arrow). Notably, this technique allowed for the deep proteomic profiling of the STB, which has been historically impossible due to the inability to derive this syncytium into a single cell suspension.

**Figure 4.**
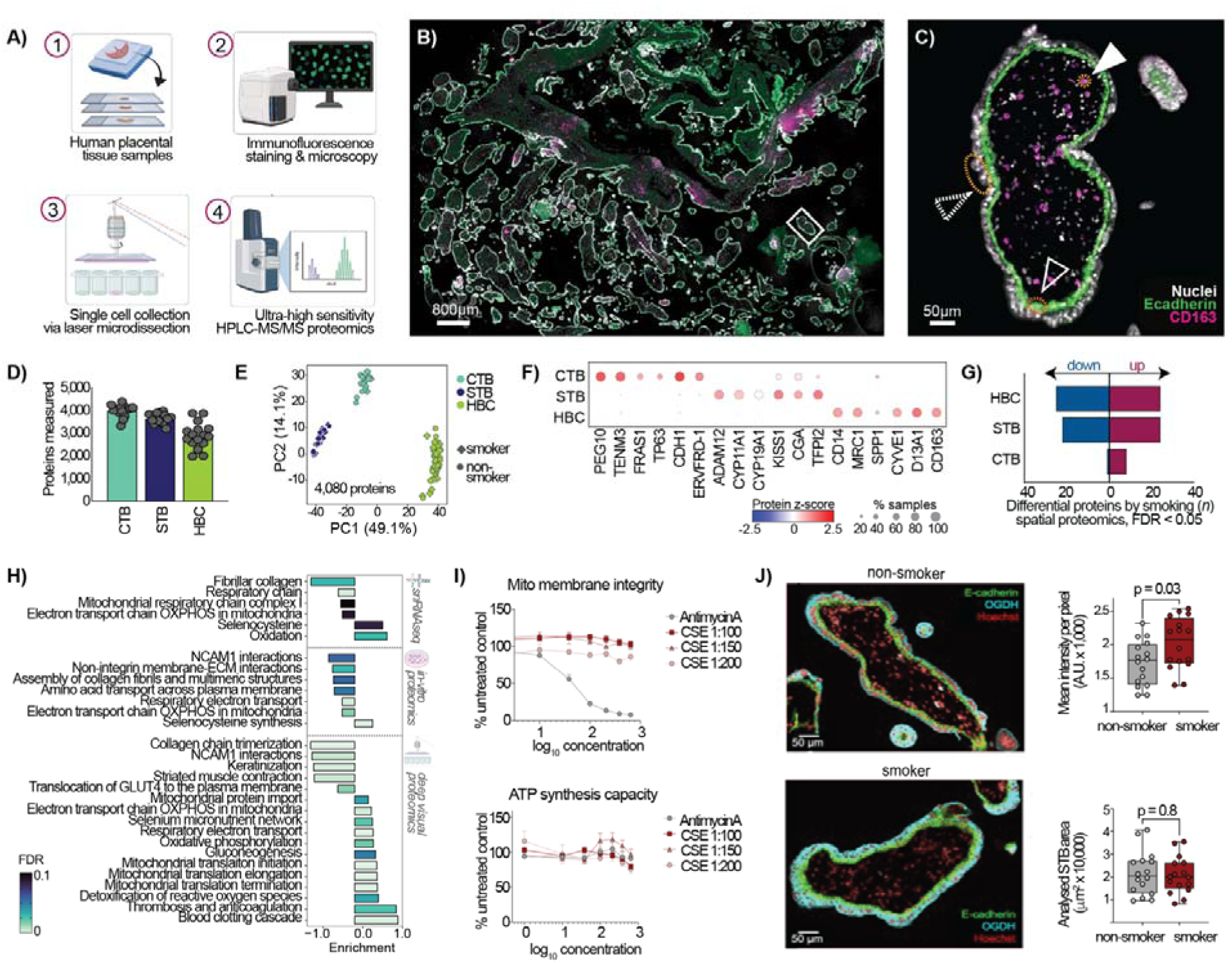
Deep visual proteomics reveals maternal smoking contributes to syncytiotrophoblast mitochondrial accumulation. **(A)** Schematic representation of the deep visual proteomics (DVP) workflow indicating four key steps. Created with biorender.com. **(B)** Representative immunofluorescence staining of first trimester control sample used for the spatial proteomics indicating cytotrophoblasts (CTB; green; E-cadherin), Hofbauer cells (HBC; magenta; CD163) and nuclei (grey; Hoechst staining of nuclear DNA). White square is indicating view field of (C). **(C)** Representative area of whole tissue immunofluorescence staining showed in B). Specific annotated regions are indicated: White arrow head, CD163+ HBC; dashed arrow head, multinucleated syncytotrophoblasts (STB); white outlined arrow head, E-cadherin+ CTB. **(D)** Numbers of proteins in microdissected CTB, STB and HBC detected by mass spectroscopy. **(E)** Principal component analysis (PCA) of proteomic profiles of microdissected CTB, STB and HBC. Each point represents a technical replicate measured (n = 3 smokers and 3 non-smokers with technical quadruplicates per cell phenotype group). **(F)** Dotplot visualizing abundance of the canonical literature-based markers in the microdissected placental cell types. Colors represent z-score of protein abundances across groups. **(G)** Number of differentially abundant tissue proteins by maternal smoking at a 5% FDR threshold. Upregulated proteins are in magenta, downregulated in blue. Differential abundance was calculated on pseudo-bulk counts per cell type or state using a generalized linear model with a Benjamini-Hochberg FDR correction. **(H)** Summary of pathway enrichment analysis of dysregulated STB from results of snRNA-Seq (upper panel), *in vitro* proteomics (middle panel) and spatial proteomics (lower panel) as indicated by scheme (absolute FC > 0.25, FDR < 0.1). Enrichment was done based on ranked log fold changes spanning the gene set enrichment analysis (GSEA), reactome and the WikiPathways databases. Positive enrichment values represent pathways enriched in smokers; negative values represent those enriched in non-smokers. Bar color is mapped to dysregulaiton FDR. **(I)** Mitochondrial toxicity analysis of *in vitro* STB (differentiated primary trophoblast stem cells (TSC)) induced by indicated concentration of cigarette smoke extract (CSE) cell culture media as percentage of untreated control media. Upper panel is presenting the mitochondrial membrane integrity, lower panel the corresponding adenosintriphosphate (ATP)-synthesis capacity. Antimycin A is representing a positive mitochondrial toxicity by inducing a reduced membrane integrity with unchanged ATP synthesis capacity. **(J)** Left panel: Representative immunofluorescence staining of first trimester villi of non-smoker (upper panel) and smoker (lower panel) indicating E cadherine (CTB; green), the mitochondrial marker 2-oxoglutarate dehydrogenase (OGDH; blue) and nuclei (red; Hoechst staining of nuclear DNA). Upper right panel: quantification of mean intensity per pixel in the STB regions analyzed using a quantitative pathology (QuPath) software. Each dot represents a measured villi area. Significance tested by two-tailed unpaired t-test with Welch’s correction. AU, arbitrary units. Lower right panel: Mean STB areas analyzed (mm2) used for the quantification of OGDH intensity per group. Significance tested by two-tailed unpaired t-test with Welch’s correction. (n = 4 smokers, n = 4 non-smokers in quadruplicate villi areas per tissue). STB, syncytiotrophoblast; CTB, cytotrophoblast; HBC, Hofbauer cell.

We microdissected and collected approximately 200 cells across the whole tissue section per phenotype technical replicate from six placental tissues (*n* = 3 smokers, *n* = 3 non-smokers, all female fetuses) also included in the snRNAseq cohort. Remarkably, we characterized phenotype-specific signatures at a detection range of 3,100 (HBC) to 4,370 (CTB) proteins (**Fig 4D**). We observed cell-type specificity of quantified samples and excellent concordance between the microdissected phenotypes of interest and their proteomic profile of canonical markers (**Fig 4E, F**). This alone constitutes the most comprehensive cell type resolved proteomic characterization of the developing human placenta to date, as this has only been previously performed at the whole-tissue lysate level at a comparable depth (4, 239 proteins)^55^.

We found vast proteomic differences in the STB and HBC between smokers and non-smokers, with a similar amount of differentially abundant (DA) proteins (FDR < 0.05; HBC, *n* = 49 DA [24 up, 25 down]; STB, *n* = 46 DA [24 up, 22 down]). The CTB displayed a remarkably low number of DA proteins, in line with snRNAseq findings (*n* = 9 DA; 8 up, 1 down) (**Fig 4G, Suppl Table 4**). STB dysregulated proteins were involved in oxidative stress response, protein folding, metabolic processes, extracellular matrix organization, immune response and pro-coagulation dynamics. Thus, we not only validated snRNAseq-based main molecular mechanisms by proteomics, but we also identified an additional role of hemostasis at the maternal-fetal interface unable to be elucidated by transcriptomics alone. In addition, we performed a deep proteomic analysis of the *in vitro* model we introduced in Fig 3 (*n* = 8,837 protein IDs). After *in-vitro* TSC differentiation towards the multinucleated endocrine STB phenotype, exposure to CSE for 48 hours led to the dysregulation of 144 proteins (FDR < 0.05) (**Suppl Table 5)**.

In the STB, we found functional pathway enrichment for selencysteine synthesis across the three modalities (snRNAseq, DVP, *in-vitro*), while markers involved in cell-adhesion, ECM integrity and cellular transport were downregulated (**Fig 4H**). Both the snRNAseq and *in-vitro* proteomics identified mitochondrial function pathways, primarily oxidative phosphorylation, to be impaired in the STB of smokers. However, in DVP results we instead found a stark upregulation of mitochondrial machinery and function related pathways (**Fig 4H**). Thus, we hypothesized that cigarette components may act as a mitochondrial toxin to the STB, inducing dysfunction and leading to accumulation in the tissue. To evaluate this, we used a cell-based multiplexed assay method that predicts mitochondrial dysfunction specifically as a result of xenobiotic insult (**Fig 4I**). We found that CSE treatment on *in-vitro* differentiated STBs had no impact on mitochondrial membrane integrity (upper panel) nor ATP synthesis capacity (lower panel), as curves deviate from our positive control mitochondrial toxin AntimycinA (**Fig 4I**). This phenomenon was consistent when cells were exposed to the same CSE concentration used for previous mRNA and proteomic experiments (1:200 dilution of 100% stock) or to higher non-lethal amounts (1:150 and 1:100 dilutions). To further reconcile these findings, we performed immunofluorescence staining of the citric acid cycle oxoglutarate dehydrogenase (OGDH) enzyme, localized to the mitochondrial membrane, on the same tissues profiled by snRNAseq and DVP (**Fig 4J**, left). We confirmed an accumulation of mitochondria in the STB of smokers compared to that of non-smokers based on STB area intensities, with comparable areas analyzed per tissue (**Fig 4J**, right; *n* = 4 areas per tissue).

Our findings shed light into the complementarity of different omics modalities in driving biological insights discovery. While soluble smoke components are not direct mitochondrial toxins to the STB, there is an adaptive accumulation of mitochondrion in the STB of smokers, likely as a response to intracellular stress and increased energy demands.

To better understand specific cell-cell interactions between the most altered cell types (STB, HBC), we used CellChat to predict dysregulated communication, integrating the results of snRNAseq and DVP proteomics results (**Fig 5A-C, Methods**). We included the CTB in these comparisons as the CTB monolayer is in direct contact with the STB and physically separates the syncytium from the HBC and stroma. To this end, we calculated paracrine interactions between the STB-CTB and the CTB-HBC as well as autocrine signaling within each cell type.

**Figure 5.**
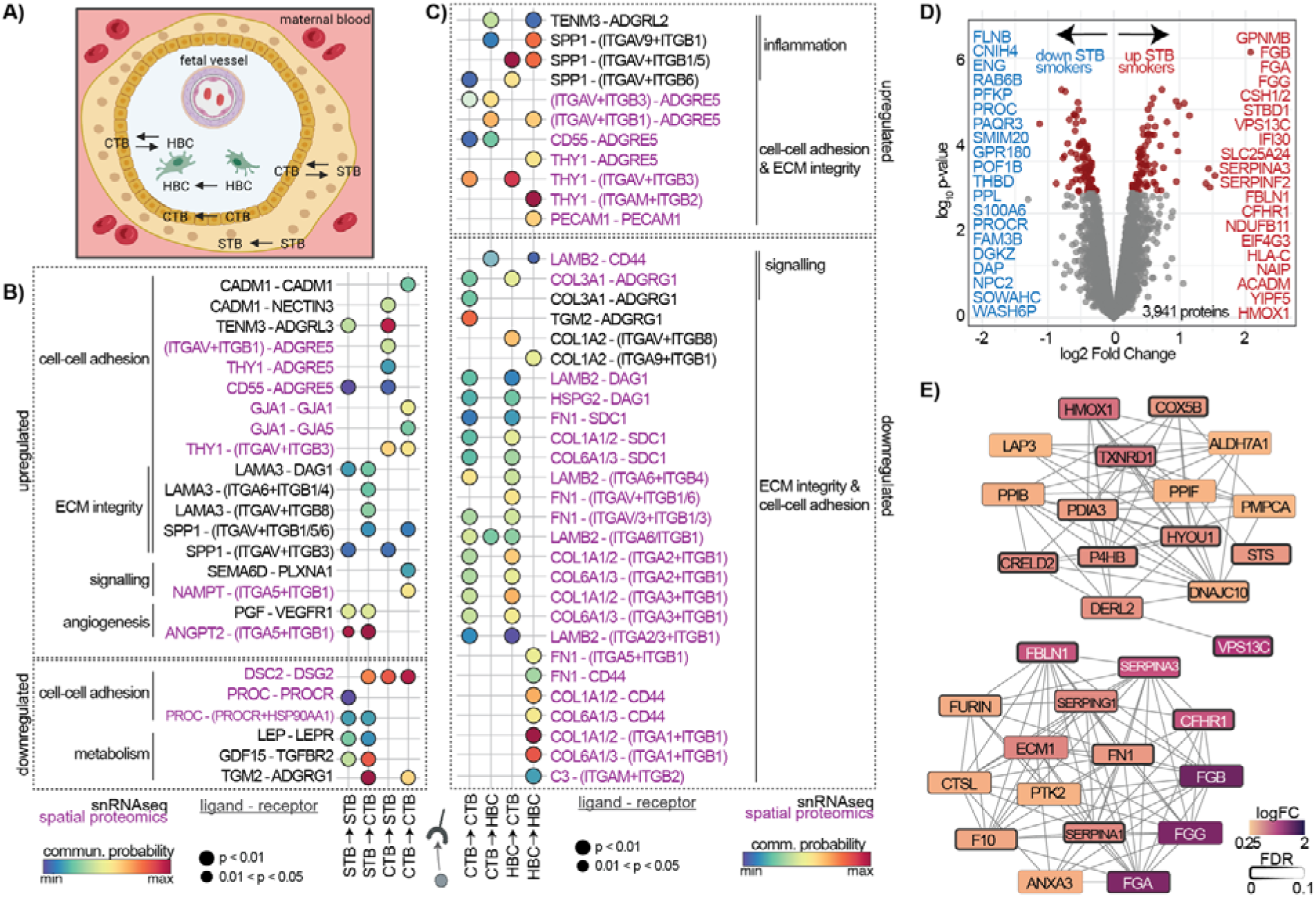
Maternal smoking impairs intra-organ communication. **(A)** Schematic representation of a placental villi cross-section bathed in maternal blood within the intervillous space. Cell-cell interactions of interest and their dysregulation are noted. Created with biorender.com. **B, C)** Significantly dysregulated cell-cell communication in the trophoblast layer **(B)** and tissue-resident immune **(C)** compartments by smoking. Ligand receptor pair communication was systematically inferred per omics modality (snRNAseq and DVP) as well as per condition separately using CellChat. Inferences per modality were integrated between conditions and interaction strength and flow compared by Welch’s t-test. Dot color indicates communication probability of ligand-receptor pairs, dot size represents the p-value, text color indicates modality of origin for significant hit. **(D)** Volcano plot of differential expressed proteins (DEGs) analysed by DVP from syncytiotrophoblast (STB). The top 20 most dysregulated genes are shown. DEGs were calculated using semi-bulk counts per cell type via a generalizer linear model and Benjamini-Hochberg correction in the edgeR package. **(E)** Hub genes of upregulated (log2FC > 0.25, FDR < 0.1) and downregulated (log2FC < -0.25, FDR < 0.1) tissue STB **(C)** and HBC **(F)** proteins by maternal smoking in early pregnancy, based on protein-protein interaction background stringDB network followed by topological analysis. Color represents log2FC, node border thickness corresponds to differential expression analysis’ FDR. **(F)** Platelets

We identified a global widespread dysregulation involving integrins, cell-adhesion transmembrane receptors acting as ECM-cytoskeletal linkers and transducers. Within the trophoblast comparison (STB-CTB), upregulated pro-angiogenic signaling was restricted to STB PGF and ANGPT2 ligands acting on the STB itself and on the CTB (**Fig 5B**). Tissue transglutaminase (TGM2) was downregulated in trophoblast communication. The *in-vitro* inhibition of this protein impairs human trophoblast intercellular fusion and disrupts stabilization of particulate material physiologically shed into the maternal circulation throughout pregnancy^56^. Further, our results suggest that dysregulated mechanical strength and stability of the trophoblast basal membrane by smoking is underpinned by the activation of the G-protein coupled receptors ADGRE5 and ADGRL3; adhesion molecules CADM1 and NECTIN3; gap-junction proteins GJA1 and GJA5; concomitant to impaired desmosome interactions (DSC2-DSG2) (**Fig 5B**). The desmosome connects CTBs to each other laterally and contributes to the attachment of the CTB to the overlying STB^57^. Within the tropho-immune comparison (CTB-HBC), we identified osteopontin (SPP1)-integrin signaling as the main driver of the pro-inflammatory HBC response (**Fig 5C**). Within the HBC only, ECM integrity was mainly impaired by fibronectin, laminin and collagen integrin signaling. In addition, we found PECAM1 signaling, a marker associated with HBC-mediated angiogenesis, upregulated in within-HBC interactions **(Fig 5C)**. In both STB-CTB and CTB-HBC analyzed, we observed an upregulation in THY1 ligand signaling, a marker and putative modulator of placental vascular development (**Fig 5B, C**). These results pinpoint specific ligand and receptor pairs likely to mediate the widespread pathway and network level dysregulation herein reported.

Another interesting finding from the DVP dysregulation of the STB was a tissue proteome enrichment of the thrombo-hemostatic network (thrombosis and anticoagulation, blood clotting cascade) (**Fig 4H**). Strongly upregulated proteins in this cell type include fibrinogen alpha (FGA), beta (FGB) and gamma (FGG) chains, as well as members of the SERPINE family (**Fig 5D, Suppl Table 4**). To reveal modulators behind the upregulated proteins, we performed network analysis and calculated hub genes that identified two distinct communities representing proteins associated with oxidative stress response, protein folding and metabolic processes (upper cluster) and proteins involved in coagulation and immune response (bottom cluster) (**Fig 5E**). Platelet activation at the maternal-fetal interface has been previously described to directly underpin or propagate placenta-associated pregnancy pathology^58^.

Through these analyses, we describe for the first time how maternal smoking disrupts placental cell communication and unveiled maternal platelet activation and fibrinogen deposits as a novel putative modulator of deleterious smoking-induced effects in the early human placenta.

## Discussion

This work unveils, for the first time, the phenotype-resolved profound impact that maternal smoking has on the transcriptomic and proteomic tissue profiles of the developing human placenta. We revealed effects of maternal smoking on all cell types and states of this organ, with the strongest dysregulation on the multinucleated and endocrine STB layer and tissue resident macrophages (HBC). Although not the most abundant cell types in early pregnancy, they play important roles in mediating placental function. The STB is in direct contact with maternal blood and plays a pivotal role as barrier interface with a unique endocrine capacity. The HBC are of fetal origin and play a fundamental role in placental development and maternal-fetal nutrient exchange. We describe orchestrated alterations in detoxification, metabolism and structural pathways in the STB and HBC that may compromise organ development and function, consequently affecting fetal growth. Our findings highlight the importance of phenotype-resolved molecular profiling in discerning processes underpinned by low-abundance yet highly physiologically relevant cell types.

Maternal smoking has been associated with sharp increases in third trimester whole-tissue oxidative stress. Hoch et al. reported decreases in ROS-mediated DNA damage (41%) in whole tissue lysates, suggesting a compensatory response to smoking-induced intracellular stress. While studies that directly investigate the impact of smoking on HBCs are sparse, Sbrana et al. described by immunostaining that at term, HBCs in placentas from smokers exhibit significantly increased oxidative DNA damage, suggesting that maternal smoking induces oxidative stress directly within fetal macrophages^59^. In our work we were able deeply phenotype the effects of smoking on the HBC specifically. We found a smoking-induced upregulation of CD14, CD63 and CD83 that indicates an activation of phagocytosis and response to placental inflammation that is also observed in chorioamnionitis^60^. A further indicator of placental inflammation was CD68 upregulation in HBC from smoking women. In a mice model for FGR, an induction by uric acid leads to an upregulation of placental inflammation and CD68^+^ macrophages^61^. Bezemer et al. proposed a putative vicious cycle in FGR where failed-maternal fetal tolerance, placental maldevelopment, oxidative stress and a placental inflammatory immune response promote maladaptive processes^5^. The high transcript and protein turnover in the HBC coupled with its pro-inflammatory profile reveal that maternal smoking conforms to this auto-amplifying vicious cycle that ultimately results in placental insufficiency.

For the first time, we provide pathway and network level evidence that intracellular oxidative stress in the trophoblast is restricted to the STB and already begins in the first trimester of pregnancy, persisting until birth^59,62^. We found no significant dysregulation in gene expression or protein abundance of hallmark markers for hypoxia, ferroptosis or apoptosis, suggesting these processes do not modulate smoke-induced STB stress. Moreover, we found no dysregulation in superoxide dismutase nor glutathione peroxidase activity, the primary free radical detoxification modulators in the human placenta at term. Instead, we discovered three main mechanisms explaining maternal smoking induced STB stress.

First, the sharp activation of downstream effectors CYP1B1 and CYP1A1 of the xenobiotic response aryl-hydrocarbon receptor pathway. The absence of free radical scavengers in maternal smokers found by us and others^63^ indicate an accumulation of active phase I enzyme metabolites unable to be adequately inactivated by phase II enzymes; contributing to oxidative stress and impaired cell function at a critical stage of placental development. Interestingly, in the intestinal epithelium, CYP1A1 and CYP1B1 activity is protective against chemical-induced damage of cell-cell tight junctions^64^. Thus, AhR activation may be protective to the CTB monolayer and explain the lower dysregulation in this cell type. Second, our findings reveal an adaptive mitochondrial accumulation within the STB of smokers, representing an early-stage energy metabolism adaptation. While this may preserve cellular function in the acute state, chronic effects pose as long-term risks to placental efficiency and fetal development. Notably, soluble cigarette smoke components did not exhibit mitochondrial toxicity *in-vitro*, despite tissue snRNAseq and *in-vitro* CSE-stimulated STB consistently indicating impaired oxidative phosphorylation (OXPHOS) capacity by cigarette smoke components. Physiological levels of accumulated mitochondrial-generated ROS likely compound to increases in intracellular stress. Third, fibrinogen deposits on the STB surface as a response to maternal coagulation cascade likely contribute to increased stressed, since platelet activation strongly triggers oxidative stress and ROS production^65^. In the healthy human placenta, STB degeneration is routinely filled by fibrin spots as a result of clotting, with up to 7% of the syncytium containing fibrin spots in the healthy placenta at term^66^. In placenta-associated pregnancy pathologies, activation of maternal platelets and increased fibrin deposition within the STB barrier either directly underpins or propagates the underlying pathophysiology^58^.

In addition, we describe widespread dysregulation of key ion, nutrient and hormone transporters in the STB, coupled with higher metabolic needs and glucose consumption. Increases in fibrin plaques can additionally alter the diffusion transport capacity of the placenta^16^. Therefore, effects of maternal smoking on the functional syncytial feto-maternal barrier can be interpreted to directly impair transport to and from the fetal compartment, adversely impacting fetal development and growth causally linked to this behavior^28^.

Furthermore, our results support a moderate preferential fusion and differentiation towards the STB phenotype in smokers. In the human placenta, CTB fusion is mechanistically governed by increases in intracellular cyclic adenosine 3’/ 5’-monophosphate (cAMP) and protein kinase A (PKA) activity induced by the autocrine-paracrine loop binding of human chorionic gonadotropin (hCG, coded by *CGA* and *CGB* genes) to the luteinizing hormone/choriogonadotropin transmembrane receptor (LHCGR). The primary modulators of this process include P300 and GCM1. Whilst we found no differences in hCG and LH levels between groups, we revealed kisspeptin (KISS1), a snRNAseq STB transition gene that directly activates cAMP pathway activity^67^ is highly upregulated in the tissue STB proteome by smoking. Additionally, in primary cell stimulation with CSE the vasoactive intestinal peptide receptor 1 protein was markedly increased. This protein directly activates adenylate cyclase (AC), the only enzyme known to generate cAMP from ATP^68^. Our findings indicate that STB fusion efficiency is indirectly affected by biochemical activator dysregulation, not directly via aberrances in morphological fusion drivers.

The inducible isoform of heme oxygenase (HO-1 encoded by *HMOX1*) is increased at the tissue protein and *in-*vitro CSE stimulated STBs gene expression and protein levels. Network analysis identified HMOX1 as a master modulator of the upregulated STB protein response to maternal smoking. HO-1 is induced in response to ROS-associated cellular stress in many tissues with protective effects^69^. The HO system also represents the main source of endogenously produced CO in the body^70,71^. CO exposure is pro-angiogenic at the murine maternal-fetal interface, without detrimental effects on pregnancy or fetal outcomes^72,73^. In human endothelial cells, iron-mediated-HO-1 activity directly regulates PGF expression and secretion^74^. Thus, we propose HO-1 induction to have a dual effect in the placental response to maternal smoking, both by contributing to oxidative stress detoxification mechanisms and by modulating increased STB pro-angiogenic *PGF* gene expression and PlGF protein secretion. *PGF* is a hub gene in the pro-angiogenesis associated cluster of upregulated STB tissue genes. This early onset phenomenon could very well modulate adaptive placental angiogenesis seen in smokers later in pregnancy^35^ that translates to decreases in placental villous blood flow and increases in blood flow resistance of the umbilical cord, leading to impaired oxygen and nutrient transport.

The relevance of CO and PlGF are further supported by the epidemiology of smoking in hypertensive disorders of pregnancy. Maternal smoking constitutes the sole environmental exposure known to reduce the risk of gestational hypertension and preeclampsia (PE) by up to 50%^75^. PE and gestational hypertension are also associated with abnormally low levels of circulating PlGF compared to those of a healthy pregnancy^76,77^, whilst we revealed an upregulation of PlGF expression and protein levels by smoking. Studies have also shown decreased HO gene expression in early pregnancy among women destined to develop PE^78^ and decreased exhaled CO among those with gestational hypertension and PE^79^. CO also induces vasorelaxation of placental vessels that decreases perfusion pressure in pregnancy^80^, which may lead to placental insufficiency by decreasing diffusion rates. We further identified several factors that play an important role in the development of PE and were inversely regulated by smoking. The already mentioned STB transition gene KISS1 (upregulated by smoking) shows lower second-trimester circulating levels in women who later on developed PE^81^. Additionally, while we detected elevated ADM expression by smoking, circulating and placental ADM mRNA have shown to be significantly reduced in women 10-12 weeks prior to PE onset^82^. A recent mice model confirmed this finding by rescuing the PE-like phenotype by nanoparticle-based forced ADM expression, and suggested this as possible intervention for PE^83^. Altogether this supports a future focus on the CO-HO-mediated trophoblast effects in the placenta to further understand PE and its largely unknown etiology to identify new therapeutic targets.

Overall, our work describes how the *in-vivo* tissue molecular landscape of the developing human placenta responds to heavy smoking in pregnancy. Smoking primarily affects the STB trophoblast syncytium and HBC, tissue-resident placental macrophages, at the transcriptomic and proteomic levels. Dysregulation at the transcriptomic and proteomic levels in STB include increases in oxidative stress, activation of the xenobiotic detoxification aryl-hydrocarbon receptor pathway and disturbed mitochondrial energy metabolism capacity. The blood clotting cascade and HO system emerged as crucial players in oxidative stress response and angiogenesis. Similar findings were observed for the HBC. Oxidative stress led to an induction of inflammation and protein translation machinery. These early molecular dysregulations have the potential to be used as a model for the manifestation of FGR later in pregnancy. In turn, giving way to explore diagnostic and therapeutic strategies to identify and mitigate identified mechanisms early in pregnancy to protect fetal health.

## Resource availability

Processed data objects used in this work including scripts for reproducing analyses and visualizations are available via GitHub (XX). Anonymized raw transcriptomic data has been uploaded to EGA under accession number XX. Anonymized raw proteomic data has been uploaded to PRIDE under accession number XX.

Additional data are provided in figures and tables in the Supplemental Material.

## Acknowledgements

We gratefully appreciate the excellent technical assistance of May-Britt Köhler, Gabriele N’Diaye, Jana Czychi, Kornelia Buttke, Aylina Deter (ECRC, MDC, Charité, Berlin) and Lena Neuper (Med. Uni. Graz). We also thank Dr Andreas Glasner (Femina-Med) for the patient recruitment of first trimester placental material.

F. Herse. and D.N.Mueller. were supported by Deutsche Forschungsgemeinschaft (HE6249/5-1; HE6249/7-1; HE6249/7-2; HE 6249/5-3; D.N.M.: Projektnummer 394046635 - SFB 1365) and BIH Omics platform. F.Herse. was funded by VolkswagenStiftung (9D289). D.N.Mueller. was supported by grants from the German Centre for Cardiovascular Research (DZHK; BER 1.1 VD). M.Gauster. was supported by the Austrian Science Fund (FWF): 10.55776/P35118, 10.55776/I6907, and 10.55776/PAT4258724. O. N. was supported through the PhD program Inflammatory Disorders in Pregnancy (DP-IDP) by the Austrian Science Fund (FWF): Doc 31-B26, PhD program Molecular Medicine at the Medical University of Graz, and through the Marietta Blau Grant (OeAD; BMBWF), and with grants from the MeFo Graz (PS-Stipendium 2019/2020). J.N. and F.C. acknowledge funding support by the Federal Ministry of Education and Research (BMBF), as part of the National Research Initiatives for Mass Spectrometry in Systems Medicine, under grant agreement No. 161L0222. F.C. received funding from the European Research Council (ERC) under the European Union’s Horizon 2020 research and innovation program (grant agreement No. 101115681) and support by the ERC (ERC starting grant). Schematic drawings were created with biorender.com.

## Author contributions

Conceptualization, D.S.V., F.H., M.G., O.N.

Resources: D.S.V., M.G., S.H., O.N.

Experiments, D.S.V., J.U., D.F., M.R., J.N.

Data analysis, D.S.V., M.R., J.N. M.P., F.C.

Figures: D.S.V., F.H.

Supervision: D.S.V., F.H., D.N.M., R.D., F.C.

Funding acquisition, F. H., M.G., O.N., F.C.

Writing the original draft: D.S.V. All authors reviewed and edited the manuscript.

## Methods

### 3.1. First trimester placenta tissue sourcing

Placental tissue was collected surgically from electively terminated pregnancies with informed consent from healthy individuals (gestational age 5 – 11 weeks). Exclusion criteria were maternal age under 18 years, maternal BMI >25 kg/m^2^ and any existing maternal pathologies. Ethical approval was obtained from the Medical University of Graz Ethics Committee (31-019 ex 18/19). Immediately after surgical extraction, tissue was stored at 4°C in culture medium (DMEM/F12 1:1, 1 g/dL glucose) and processed in less than 4 hours. Villous tissue was rinsed twice in cold (4°C) 0.9% NaCl solution to remove blood, and either snap frozen in liquid nitrogen and stored at -70°C until processing; or fixed with 10% formalin, paraffin embedded and dehydrated according to standard protocols. Table of participant characteristics is presented in **Table 1**. Values were tested for normality via Shapiro-Wilk and Anderson-Darling tests and compared without removing outliers using either two-tailed unpaired Welch’s t-test or two-tailed Mann-Whitney U test.

### 3.2. Sample processing

#### 3.2.1. snRNAseq

Approximately 100-200 mg frozen placental tissue was processed according to an optimised nuclei isolation protocol by Krishnaswami et al^134^. Briefly, frozen tissue was crushed in liquid nitrogen using a frozen mortar and pestle and disrupted with a pre-cooled glass Dounce homogenizer in homogenisation buffer (1X NIM2 [1X protease inhibitor, 1 μM DDT, 250 mM sucrose, 25 mM KCl, 5 mM MgCl2, 10 mM pH8.0 Tris], 0.4 U/µL RNAseInhibitor, 0.2 U/µL SUPERase-in, 0.1% v/v Triton X-100) and filtered through a FACS tube with a 35 µm sieve cap. Homogenate was incubated in the dark, on ice, for two minutes with DAPI (5 µg/µL) and centrifuged for eight minutes (1,000 RCF, 4°C). Pellets were resuspended with staining buffer (3% BSA RNase free + 1% SUPERase-in in DPBS), transferred to a FACS-tube with a 35 µm sieve cap and analysed using the BD FACSAria III cell sorter using the BD FACSDiva software (v6.1.3). After FACS sorting with a 90% viable single nuclei cut-off, nuclei in landing buffer (DPBS with 4% BSA RNAse free + 10% RNAseInhibitor + 10% SUPERase-in) were counted using a digital counting chamber to 400-500 nuclei/µl and loaded onto 10x Genomics Chromium chips. Single-index v3 libraries were prepared according to manufacturer’s instructions (Chromium Single Cell 3’ Kits v3.1 Dual Index User Guide – CG000315). Libraries were sequenced on an Illumina HiSeq-4000 (pair-ended) with a minimum coverage of 50,000 raw reads per nucleus.

#### 3.2.2. Primary cell culture

##### Trophoblast stem cells

Villous cytotrophoblasts (vCTBs) were isolated from first trimester human villi according to published protocols^135^. Precisely, placental tissue (6^th^ – 8^th^ week of gestation) was cut from chorionic membranes, minced into small pieces and digested thrice for 10 min, 15 min and 15min (HBSS, 0.25% Trypsin, 1.25 mg/mL DNAse I; 5mL solution per mL tissue). Each digestion was stopped using 10% FBS [v/v]. Cells were filtered through a 100-μm pore size cell strainer, and cells from the second and third digestions were pooled. Cells were loaded onto Percoll gradients (10 – 70 % [v/v]) and vCTBs were collected between 35 – 50 % of Percoll layers, pelleted, and washed twice with HBSS. Red blood cells were removed by incubating with erythrocyte lysis buffer (155 mM NH_4_Cl, 10 mM KHCO_3_, 0.2 mM EDTA, pH 7.3) at RT for 5 min. vCTBs were washed twice with HBSS and seeded at a density of 0.5 x 10^6^ cells per well of a 6w plate pre-coated with 5 µg/mL collagen IV. Cells were cultured with 2mL/well of published trophoblast stem cell (TSC) media (**Table 2**) ^49^at 37°C and 5% CO_2_, media was replaced every second day^16^. When reaching 70-80% confluency cells were enzymatically dissociated with 1mL TrypLE for 10 minutes at 37°C and passaged at a 1:2 ratio. Following the 10^th^ passage, only proliferative cells with a TSC phenotype remain, which were passaged as above at a 1:3 split ratio every three days and used for differentiation and stimulation experiments between passages 15 and 25. Cells were routinely frozen in cell banker 2 (2 – 5 x 10^6^ cells per mL) for cryogenic storage.

**Table 2.** Culture medium recipes used for *in-vitro* culture work.

| Cell type | Culture medium |
| --- | --- |
| TSC | DMEM/F12 Glutamax, supplemented with: 0.3 % BSA, 5 ng/mL Gentamycin, 0.1mM 2-ME, 1 % ITS-X, 0.2 % FBS, 5 μM ROCKi, 2 μM CHIR99021, 0.5 μM A83-01, 0.8 mM VPA and 50 ng/mL hEGF. |
| STB | DMEM/F12 Glutamax, supplemented with: 0.3% BSA, 5 ng/mL Gentamycin, 0.1 mM 2-ME, 1 % ITS-X, 2 μM Forskolin and 2.5 μM ROCKi. |

Concentrations reported are final concentrations used in supplemented medium. TSC, trophoblast stem cell; STB, syncytiotrophoblast; BSA, bovine serum albumin; 2-ME, beta Mercaptoethanol; ITS-X, insulin-transferrin-selenium-ethanolamine; FBS, fetal bovine serum; ROCKi, rho-associated protein kinase inhibitor; CHIR99021, glycogen synthase
kinase 3 (GSK-3) inhibitor; A83-01, activin/NODAL/TGF-beta pathway inhibitor; VPA, valproic acid sodium salt; hEGF, human epidermal growth factor.

Following 24 hours attachment in TSC medium (**Table 2**), TSCs were stimulated with CSE diluted 1:200 in supplemented medium for up to 24 hours. At the hour (h) timepoints 3 h, 6 h, 9 h, 12 h and 24 h, 700 μL QIAzol reagent was added directly to each well and frozen at - 70°C prior to RNA isolation and cDNA synthesis. To differentiate towards the STB lineage, TSCs were harvested upon reaching ∼80 % confluency with TrypLE for 10 minutes at 37°C and seeded at a density of 0.1 x 10^6^ cells per well of 12w plate pre-coated with 2.5 μg/mL collagen IV. After attachment for 24 hours, cells were cultured in 1mL STB medium (**Table 2**). Medium was replaced daily for the first four days. On day 6, cells were stimulated for 6 hours with CSE 1:200 in STB medium and 700 μL QIAzol reagent was added directly to the wells and frozen at -70°C prior to RNA isolation and cDNA synthesis. Methods for the STB differentiation and processing of pellets analyzed by LC-MS are outlined in the proteomics section below (section 3.2.6).

#### 3.2.3 Cigarette smoke extract

##### Cigarette smoke extract (CSE) generation

The reproducible collection of nicotine, cotinine and aqueous constituents of cigarette smoke in the form of CSE was performed using an established protocol with some modifications^54^. The apparatus to collect aqueous components was built without modifications. In short, the cap of a 50 mL conical tube was cut twice, with a quick-connect coupler inserted to one hole and 1.5 cm of 40 cm of small tubing to the other. Epoxy resin was applied to the cap using a cotton tip and let to set overnight. The next morning, 25 cm of large tubing was attached to the coupler on one side, and a 1 mL pipette tip to the other side. The pipette tip was replaced with each batch whilst tubing remained untouched. The cap screwed onto the conical tube containing 35 mL of basal DMEM/F12 GlutamaX medium used for TSC and STB cultures (**Table 3**). Inside a chemical hood, the apparatus was fastened to a ring stand with 2 three-prong clamps. A 50mL glass syringe was connected to the small tubing and a reference 1R4F cigarette to the large tubing. The cigarette was marked at 23 mm including the filter. The syringe was greased with a single layer of petroleum jelly. To produce CSE, short ‘puffs’ (syringe drawn to 35 mL in 2-4 sec) followed by a 30 sec delay were performed. During this delay, the syringe was disconnected from the apparatus and smoke ‘exhaled’ by depressing plunger onto a paper towel. Through this, mainstream cigarette smoke was bubbled through the cell culture medium and soluble components captured to model human smoking. This process was repeated until reaching the 23 mm mark and number of ‘puffs’ recorded per cigarette, denoting a technical batch replicate. Per bacth replicate, 13 cigarettes were used. Four 100% CSE batches with 10 pooled technical replicates in each were made. Pooled batches were adjusted to 7.4 pH by adding 1M NaOH, sterile filtered through a 0.2 μM mesh, aliquoted and frozen at -70°C for characterization and use in *in-vitro* stimulations.

**Table 3.**
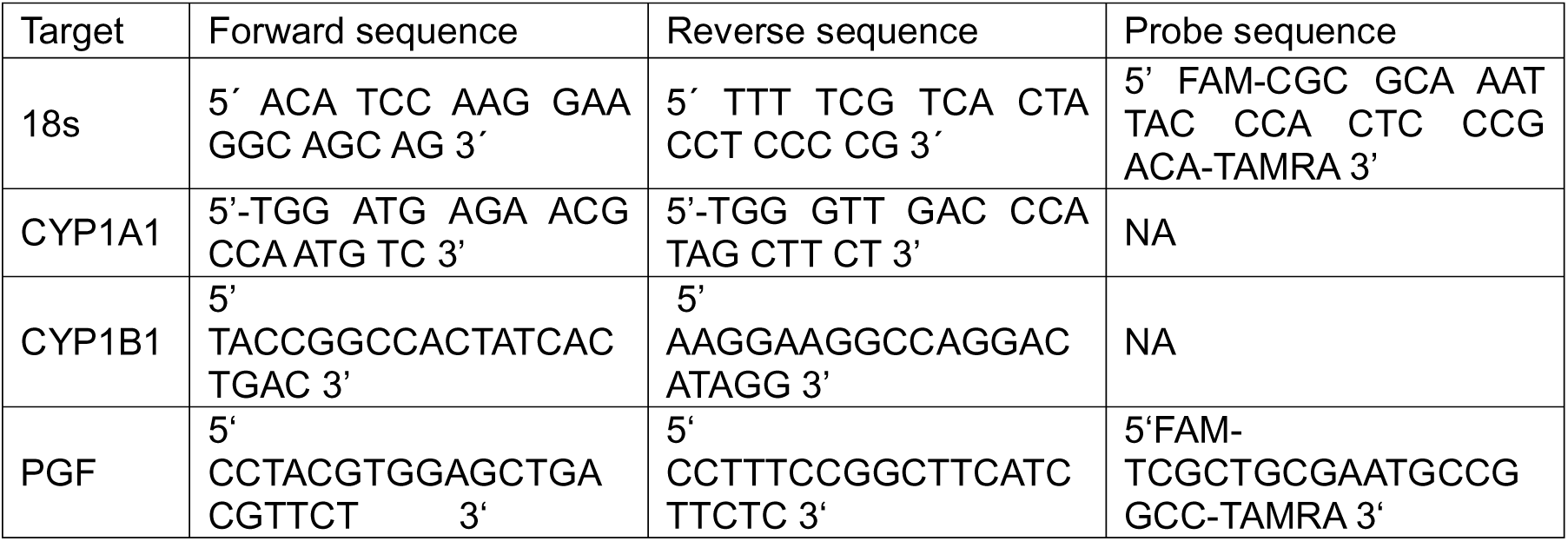
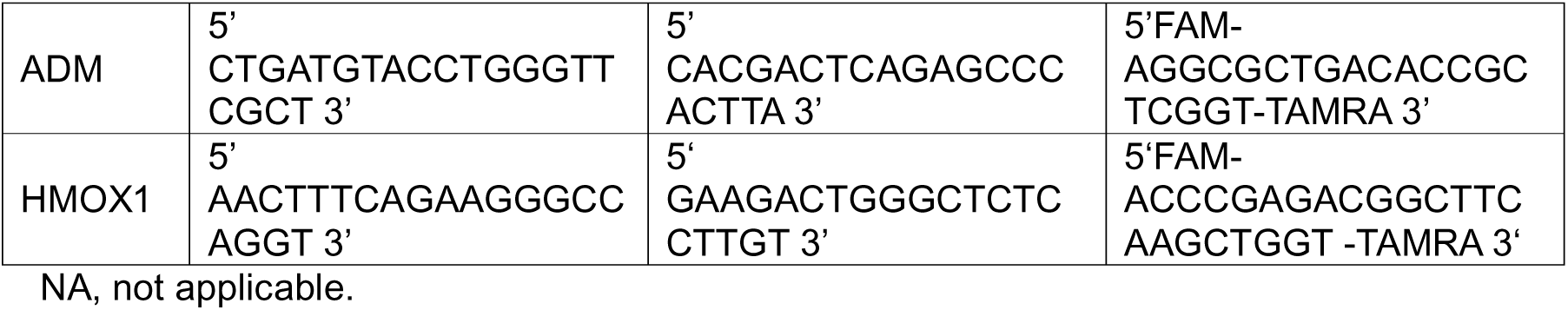
Oligonucleotide primer sequences used to evaluate mRNA expression in independent validation cohort.

##### Nicotine and cotinine quantification in CSE

Quantification of nicotine and cotinine was done as described in Shin et al^84^. In short, A 20-mL glass test tube was used to hold 5 mL of CSE. Approximately 300 mg of potassium carbonate (K_2_CO_3_) and 50 μL of diphenylamine as an internal standard were added to the solution, which was then extracted with 7 mL of ethyl ether through 10 minutes of mechanical shaking. The organic phase was transferred to another 20-mL test tube containing 20 μL of acetic acid and dried using a nitrogen stream to a volume of 50 μL. The solution was further dried with around 100 mg of sodium sulfate before a 2-μL sample was automatically injected into the gas chromatography system. Calibration curves for nicotine and cotinine were established by extraction after adding standard amounts ranging from 1.0 to 5000 ng and 0.5 μg of internal standard to 5 mL of CSE. The ratio of the peak area of the standard to that of the internal standard was used to quantify the analytes. Mass spectra were obtained using an Agilent 6890/5973 N instrument, with the ion source operating in electron ionization mode at 70 eV. Full-scan mass spectra (m/z 40–800) were recorded for analyte identification. Separation was achieved using an HP fused-silica capillary column with cross-linked methylsiloxane. Samples were injected in split mode with a splitting ratio of 1:8, and the helium flow rate was 1.0 mL/min. Operating parameters include: injector temperature, 280°C; transfer line temperature, 300°C; and oven temperature, programmed from 80°C at 20°C/min to 300 °C (held for 5 min). The ions selected were m/z 84, 133, and 161 for nicotine, and m/z 98 and 176 for cotinine, and m/z 168 and 169 for diphenylamine.

##### VOC Screening of CSE by means of GC-MS

Volatile organic compounds (VOCs) were automatically injected into the GC using an Agilent 7697A static headspace sampler (SHS, Agilent Technologies, Vienna, Austria). For this purpose, 3.5 mL of 100% pooled CSE stock medium was transferred to a 20 mL headspace vial and spiked with nicotine-d_4_ and 2-fluorobiphenyl as internal standards to achieve a final concentration of 0.99 and 1.00 µg per vial, respectively. To facilitate evaporation of VOCs in the headspace of the vial, a pre-equilibration temperature of 95°C was applied for 15 min before the analytes were automatically transferred to the GC inlet operated at 250°C. A 1 mL SHS loop was selected and maintained at 140°C with a transfer line temperature of 155°C to prevent compound loss. Helium (purity > 99.999%) was selected as GC carrier gas at a flow rate of 1.8 mL/min with a split ratio of 1:5. The separation was performed on an Agilent 5977B gas chromatograph using a cross-linked Agilent DB-Wax column with 30 m x 250 µm I.D. x 0.50 µm film thickness. The GC oven was initially held at 45°C for 2 min, followed by a thermal increase of 15°C/min to 260°C, held for 2 min. Detection was performed on an Agilent 5977 single quadrupole mass spectrometer (Santa Clara, CA, USA) operated in EI mode at 70 eV and a source temperature of 230°C. The single quadrupole was operated at 150°C in SCAN mode with m/z range from 33-550. Different solvent polarities were selected for the isolation of semi-volatile organic compounds, whereby a total of 20 mL of pure CSE medium was extracted 5 times with 7 mL of methylene chloride, followed by extraction with *tert*-methyl butyl ether and hexane, respectively. In this way, a total extract volume of 35 mL was obtained for each solvent, which was concentrated to a final volume of 150 µL containing 9.14 µg/mL of 2-fluorobiphenyl as an internal standard. A total of 2 µL of the extract was injected directly into the above-mentioned GC-MS system with similar parameters except for an adapted Agilent DB-Wax column with 60 m length x 250 µm I.D. x 0.25 µm film thickness operated with an extended temperature programme set to 260 °C for up to 10 min. Medium controls were included in the sample preparation and analysis workflow for all samples subjected to SHS or direct injection GC-MS. Peak picking was performed on spectral data using Agilent Masshunter Unknown Analysis (v10.1) and manually controlled using Masshunter Quantitative Analysis (v10.1). The peaks obtained were identified based on an in-house reference library build from certified reference standards together with a Wiley-National Institute of Standards and Technology (NIST) library search, with subtraction of corresponding sample blanks and an area threshold of at least 5000 counts. The analytes were quantified from their peak areas relative to the area of the respective reference standard or the area obtained for the internal standard 2-fluorobiphenyl.

#### 3.2.4. RNA isolation and RT-qPCR

Cells from experiments previously described or pulverized tissue were lysed in QIAzol lysis reagent. RNA was isolated using the Qiagen RNease mini kit according to the manufacturer’s instructions. RNA quality was determined using an Agilent 2100 Bioanalyzer. Quality check was followed by reverse transcription of 1 μg total RNA per reaction using the Applied Biosystems High-Capacity cDNA Reverse Transcription Kit according to the manufacturer’s manual. For qPCRs, the QuantStudio 3 Real-Time PCR System with either TaqMan Fast Universal PCR Master Mix or Fast SYBR Green Master Mix were used. Primer and probes (**Table 3**) were designed using Real-time PCR (TaqMan) Primer and Probes Design Tool (online tool) from GenScript and synthesized by the company BioTez GmbH, Germany. Primers were diluted to a final concentration of 10 mM and probes to a final concentration of 5 mM. The target mRNA expression was quantitatively analysed using the standard curve method in the QuantStudio Design and Analysis Software (v2.6.0). All expression values were normalized to the housekeeping gene *18S*. Resulting values were plotted, tested for normality via Shapiro-Wilk and Anderson-Darling tests and compared between groups without removing outliers using either two-tailed unpaired Welch t-tests or two-tailed Mann-Whitney U test in GraphPad Prism software (v10.2.0).

#### 3.2.5. Immunofluorescence staining

Formalin fixed paraffin embedded (FFPE) placenta tissue sections (5 μm) were mounted on Superfrost Plus slides and dried overnight. Standard deparaffinisation was followed by antigen retrieval in the incubator with Tris/EDTA pH9 solution for 20 min at 93°C, cooled for 20 min, transferred to warm distilled water for 5 min and cooled for 5 min. Thereafter, sections were washed with PBST (PBS + 0.1% Tween 20) and blocked by incubation with Ultra V Block for 10 min at RT. For OGDH & mouse E-cadherin double staining, primary antibodies were diluted in primary antibody solution: PBST + 1% normal goat serum (NGS) + primary antibodies (**Table 4**) and incubated on sections overnight at 4°C. The following morning, slides were washed three times with PBST, stained with secondary antibody solution: PBST + 1% NGS + mouse IgG-Cy3 and rabbit IgG-AF488 (**Table 5**) and incubated in the dark at RT for 1 hour. After secondary antibody staining, slides were washed three times with PBST and mounted with VectaShield medium with DAPI. Rabbit immunoglobulin fraction and negative control mouse IgG were used as described above and revealed no staining. Slides were imaged using the Zeiss Axioscan 7 slide scanner. Regions of interest representing STB areas from four villi per tissue sample were selected using the open-source software for digital pathology image analysis QuPath (v0.4.2) and intensity per pixel quantified in a blinded fashion.

**Table 4.**
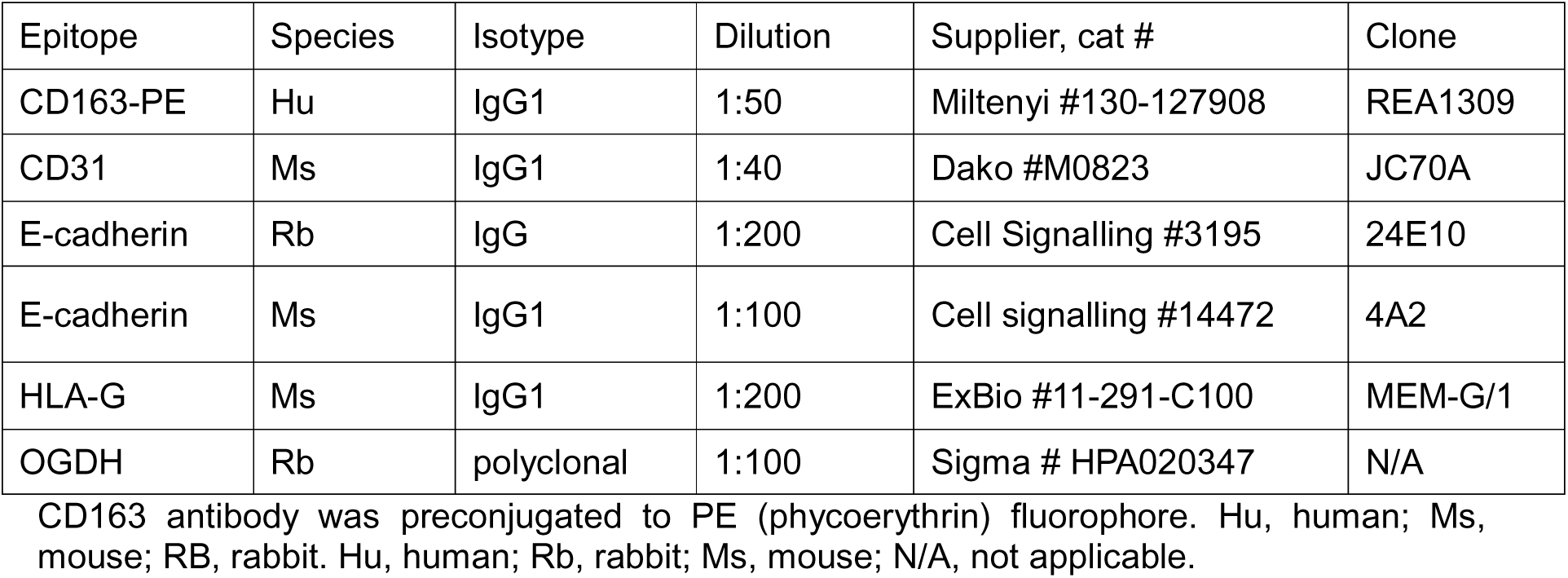
Primary antibodies used for FFPE tissue immunofluorescent staining.

| Epitope | Species | Isotype | Dilution | Supplier, cat # | Clone |
| --- | --- | --- | --- | --- | --- |
| CD163-PE | Hu | IgG1 | 1:50 | Miltenyi #130-127908 | REA1309 |
| CD31 | Ms | IgG1 | 1:40 | Dako #M0823 | JC70A |
| E-cadherin | Rb | IgG | 1:200 | Cell Signalling #3195 | 24E10 |
| E-cadherin | Ms | IgG1 | 1:100 | Cell signalling #14472 | 4A2 |
| HLA-G | Ms | IgG1 | 1:200 | ExBio #11-291-C100 | MEM-G/1 |
| OGDH | Rb | polyclonal | 1:100 | Sigma # HPA020347 | N/A |

CD163 antibody was preconjugated to PE (phycoerythrin) fluorophore. Hu, human; Ms, mouse; RB, rabbit. Hu, human; Rb, rabbit; Ms, mouse; N/A, not applicable.

**Table 5.** Primary antibodies used for FFPE tissue immunofluorescent staining.

| Isotype specificity | Host | Fluorophore | Dilution | Supplier, cat # |
| --- | --- | --- | --- | --- |
| Ms IgG | Goat | AF-647 | 1:1000 | Invitrogen, A21235 |
| Rb IgG | Goat | AF-488 | 1:1000 | Invitrogen, A11008 |
| Ms IgG | Goat | Cy3 | 1:1000 | Jackson Laboratories, #715-165-151 |

Ms, mouse; Rb, rabbit.

#### 3.2.6. Proteomics

##### Deep visual proteomics (DVP)

Formalin fixed, paraffin embedded (FFPE) placenta tissue sections (5 μm) were mounted on PPS FrameSlides (cat.no. 11600294) and dried overnight. Standard deparaffinisation was followed by antigen retrieval in the incubator with Tris/EDTA pH9 solution for 20 min at 93°C, cooled for 20 min, transferred to warm distilled water for 5 min and cooled for 5 min. Thereafter, sections were washed with PBST (PBS + 0.1% Tween 20) and blocked by incubation with Ultra V Block for 20 min at RT. For triple staining (CD163, CD31, E-cadherin), primary antibodies were mixed and diluted in primary antibody solution: PBST + 1% normal goat serum (NGS) + primary antibodies (**Table 4**) and incubated on sections overnight at 4°C. The following morning, slides were washed three times with PBST, stained with secondary antibody solution: PBST + 1% NGS + mouse IgG-AF647 & rabbit IgG-AF488 (**Table 5**) and incubated in the dark at RT for 1 hour. After secondary antibody staining, slides were washed three times with PBST and mounted with Slow Fade Diamond Mounting Medium (cat.no. 36964) with DAPI. Serial tissue sections were stained with HLA-G to exclude the selection of cell column trophoblasts as regions of interest. Rabbit immunoglobulin fraction and negative control mouse IgG were used as described above and revealed no staining. Slides were imaged using the Zeiss Axioscan 7 slide scanner using a 10X objective and 2×2 binning. Regions of interest representing cell phenotypes were manually labelled using the open-source software for digital pathology image analysis QuPath (v0.4.2). Regions of interest were collected by laser microdissection (LMD) on a Leica LMD7 microscope using a 63x objective operated in brightfield mode. An area of approximately 50,000 µm^2^ was collected in quadruplicate technical replicates per phenotype per biological sample into individual wells of a 384-well plate.

Collected cells were processed for bottom-up LC-MS based proteomics as recently described^45^, with small adjustments. Briefly, acetonitrile (2×10uL) was pipetted to each well to drag samples to the bottom of their wells, and later evaporated with rotovap. For cell lysis, 4 µl of 60 mM triethylammonium bicarbonate (TEAB) was added to each well, shortly centrifuged (2,000 RCF, 1 min) and the plate heated at 95°C for 60 min in a thermal cycler (Bio-Rad, 384-well reaction module) at a constant lid temperature of 110°C. 1 µl of acetonitrile (ACN) was then added to each well (20% final concentration) and heated again at 75°C for 60 min in the thermal cycler. Samples were shortly cooled to room temperature and 2 µl LysC added pre-diluted in ultra-pure water to 2 ng/µl and digested for 4 h at 37°C in the thermal cycler. Subsequently, 2 µl trypsin (Promega Trypsin Gold) was added pre-diluted in ultra-pure water to 2 ng/µl and incubated overnight at 37°C in the thermal cycler. The next day, digestion was stopped by adding trifluoroacetic acid (TFA, final concentration 1% v/v) and samples vacuum-dried (approx. 60min at 60°C). Samples were stored at -20°C until LC-MS availability. Finally, 4 µl MS loading buffer (3% ACN in 0.2% TFA) was added, the plate vortexed for 10s and centrifuged for 5 min at 2,000 RCF.

##### STB pellet proteomics sample preparation

Primary human trophoblast stem cells (TSCs) cultured in TSC medium (**Table 2**) were harvested upon reaching ∼80% confluency with TryplE for 10 minutes at 37°C and seeded at a density of 0.1 x 10^6^ cells per well of a 12w plate pre-coated with 2.5 μg/mL collagen IV. After attachment for 24 hours, cells were cultured in STB medium (**Table 2**) to direct differentiate them towards the STB lineage^49^. Medium was replaced daily for the first four days. Between days 5 and 7 media was replaced every 12 hours in the presence or absence of cigarette smoke extract (CSE) diluted 1:200 in STB medium. At days 0, 5, 6 and 7 cells were harvested with TryplE for 12 minutes at 37°C and centrifuged for 6 min at 300 RCF to collect pellets. Pellets were transferred to 1.5 mL reaction tubes, washed thrice with PBS for 5 min at 300 RFC, air dried and frozen at -70°C. Cell pellets were lysed with 50uL of sodium deoxycholate lysis buffer (1% (w/v) SDC, 150 mM NaCl, 100 mM Tris pH 8, 1 mM EDTA, 10 mM DTT, 40 mM CAA, 1:100 phosphatase inhibitor cocktail), each tube was vortexed, spinned down, and transferred to a 96 well plate. The samples were incubated for 10min at 95°C, then 25 U of benzonase were added to each sample. Peptide digestion took place by adding 1uL of 10 ng/uL trypsin/LysC mixture (1:50 ratio), then incubating samples at 37°C overnight. 5uL of 10 % formic acid (FA) was added to stop lysis, and precipitate the SDC salts, then the supernatant for each well was transferred to a new plate, if needed 1 % FA was added to dilute samples to a suitable volume. Samples were then washed by stage-tipping using EVOSEP Evotip Pure tips with Agilents’s AssayMAP Bravo. Protein concentration was measured by nanodrop, and each sample was diluted in a new plate to 100 ng/uL.

##### Liquid chromatography mass spectrometry (LC-MS) analysis

For both DVP and cell pellet proteomics, LC-MS analysis was performed with an EASY-nLC-1200 system (Thermo Fisher Scientific) connected to a trapped ion mobility spectrometry quadrupole time-of-flight mass spectrometer (timsTOF SCP, Bruker Daltonik GmbH, Germany) with a nano-electrospray ion source (Captive spray, Bruker Daltonik GmbH). Peptides were loaded on a 20 cm in-house packed HPLC-column (75 µm inner diameter packed with 1.9 µm ReproSilPur C18-AQ silica beads, Dr. Maisch GmbH, Germany). Peptides separation followed a gradient with a flow rate of 250 nL with increasing concentration of buffer B (0.1% formic acid, 90% ACN in LC-MS grade H_2_O) to 60%. Buffer A (3% ACN, 0.1% formic acid in LC-MS grade H_2_O). The total gradient length was 21 min and 44 min, for DVP and STB pellet samples, respectively. Column temperature was controlled by a column oven and kept constant at 40°C. Mass spectrometric aquisition was performed in data-independent (diaPASEF) mode^85^ using the default method for long gradients with a cycle time of 1.8 s. Ion accumulation and ramp time in the dual TIMS analyser was set to 100 ms each and we analysed the ion mobility range from 1/K0 = 1.6 Vs cm-2 to 0.6 Vs cm-2. The total m/z range was set to 100-1,700 m/z. The collision energy was lowered linearly as a function of increasing mobility starting from 59 eV at 1/K0 = 1.6 VS cm-2 to 20 eV at 1/K0 = 0.6 Vs cm-2. Singly charged precursor ions were excluded with a polygon filter (timsControl software, Bruker Daltonik GmbH).

#### 3.2.7. Enzyme-Linked Immunosorbent Assays

The blood samples from women in the first trimester of pregnancy used were taken prior to the surgical elective termination procedure, and serum isolated by routine centrifugation protocols. Samples were analyzed for cotinine (Abnova #KA0930) and placental growth factor (R&D Systems #DPG00) concentrations according to manufacturer’s instructions. Absorbance was read at 450 nm (with 540nm wavelength correction) using the spectrometer SPECTROstar Nano. For the cotinine ELISA, assay sensitivity is 1 ng/mL, the cut-off for smokers was set at 3 ng/mL cotinine and cross reactivities include nicotine <1 %, nicotinamide <1 % and nicotinic acid <1 %. For the PlGF ELISA, assay sensitivity is 7 pg/mL, no cut-offs were used and no reported significant cross-reactivities are present.

### 3.3. Data analysis

#### 3.3.1. snRNAseq

##### Data processing and quality control

The demultiplexing, processing, identification of Unique Molecular Identifiers (UMI) and barcode filtering of raw 3’ snRNA-Seq data was performed using Cell Ranger software (v 6.1.2) from 10x Genomics. The transcripts were aligned against the pre-built human reference genome GRCh38 premRNA version 3.0.0, which was built from the GRCh38 precompiled reference, and modified for use with snRNA-Seq data by extracting “transcripts” features from the gene model GTF and instead annotating these as “exon”, as described in the protocol defined by 10x Genomics (https://support.10xgenomics.com/single-cell-gene-expression/software/release-notes/build#grch38_3.0.0). Technical systematic background noise including ambient RNA molecules, random barcode swapping from raw (UMI) matrices and empty droplets were optimised per sample and eliminated using the CellBender package (v0.2.2) in python (v3.7.0) with 18,000 total number of droplets, 150 epochs, 0.01 fpr using a combined ambient and swapping model. Technical doublets were modelled per sample using DoubletFinder (v2.0.3). Subsequently, pre-processed matrices were loaded into R (v4.1.2) and further processed using Seurat (v4.1.0). Genes expressed in fewer than 10 nuclei per sample were removed from the count matrices. Within-sample nuclei having fewer than 200 expressed genes, a log10 ratio of genes per UMI <. 0.80, more than 0.5% expression belonging to mitochondrial genes or having a single-gene expression spanning > 30% of counts were filtered out and excluded. Distributions of the abovementioned metrics were evaluated and recorded. Three sequenced tissue samples did not pass quality control expectations and were excluded from the study which included two smokers and one control samples. Ultimately, 88,808 nuclei were included with an overall capture of 28,269 genes and a mean of approximately 4,000 genes mapped to any given nucleus. Fetal sex was inferred based on gene expression patterns of female-associated (*XIST, PCDH11X, ZFX*) and male-associated (*PCDH11Y, USP9Y, DDX3Y, TTTY14*, *EIF1AY*).

##### Data integration and cell type and state annotation

Dataset integration was performed to model and correct for gene expression heterogeneity in samples attributable to differences in biological (sample ID, fetal sex) and technical (batch, sequencing depth) variation using the ‘sctransform’ package (v0.3.3). A zero-inflated negative binomial distribution was used to model gene expression. Anchor features across datasets were computed with the ‘FindintegrationAnchors’ function using the 4,000 most variable genes computed with the ‘SelectIntegrationFeatures’ function. With integration anchors as foundation, datasets were integrated after normalization and variance stabilization with the ‘SCTransform’ function without regressing mitochondrial counts or sequencing depths. Cell cycle state was annotated using cell cycle scores with the ‘CellCycleScoring’ function that predicts S, G2M and G1 phases based on expression correlation of canonical marker sets.

Gene expression counts from the integrated dataset was used for cell type and state annotation. Gene expression counts from non-integrated data were used for downstream analyses.

Linear dimensionality reduction was performed on the 4,000 most variable genes of the integrated dataset and 50 principal components (PCs) were calculated. The K-nearest neighbour graph was computed using the first 30 PCs. Uniform Manifold Approximation and Projection (UMAP) visualisation was used to further reduce the high dimensional latent spaces to 2D using the first 50 PCs. In this visualisation, dots correspond to specific nuclei with unique x,y coordinates. Louvain clusters were computed in an unsupervised manner based on the k-nearest neighbour graph at resolutions between 0.1 and 2 in 0.1 step increments using the Louvain algorithm with modularity optimizer version 1.3.0 by Ludo Waltman and Nees Jan van Eck, including a random seed for reproducibility. The most appropriate resolution (1.0 yielding 36 clusters) to use for annotation was selected based on a cluster tree relationship visualisation at all resolutions using the clustree package (v0.5.1). Differentially expressed and conserved genes between clusters was calculated using the logistic regression based ‘FindAllMarkers’ and ‘FindConservedMarkers’ functions in Seurat package, respectively. This information was used for annotation in combination with the cluster-specific expression of literature-based canonical markers. One contaminating cluster (*n* = 474 nuclei) negative for all placental canonical markers and uniquely expressing maternal decidua epithelial cell markers *PAX8, PAEP* and *CP* was removed. Ultimately, clusters with unique transcriptomic fingerprints in relation to the rest of the dataset were annotated as their own cell type or state, whilst clusters with conserved transcriptomic fingerprints were merged to belong to the same type or state.

##### Differential gene expression analysis

Differential expression was performed on raw RNA counts from all annotated cell types and states excluding proliferating erythroblasts (EBp) and fibroblasts (FBp) due to their low abundances. Counts and associated metadata from Seurat were used to create a SingleCellExperiment object (v.1.22.0). Pseudo-bulk gene expression profiles were generated by aggregating counts for nuclei of the same cell phenotype and tissue sample ID using the ‘aggregateAcrossCells’ function in the scuttle package (v1.4.0).

Differentially expressed genes (DEGs) per cell phenotype were empirically calculated using the edgeR package (v3.36.0). First, a DGEList object from the pseudo-bulk profiles was created and genes that were not expressed above a log-CPM threshold in a minimum of five samples (the size of the smallest group in our experimental design) were filtered out. Nuclei with fewer than ten counts were also removed. Normalization factors were calculated using the trimmed mean of M-values (TMM) method to account for composition biases in the data. A design matrix was constructed and included batch and fetal sex to model the effects of these variables on gene expression. Robust dispersion estimates were obtained using the ‘estimateGLMRobustDisp’ function, which accommodates for potential outliers. A negative binomial generalized linear model was fitted genewise using the ‘glmFit’ function with robust parameter settings to compute the maximum likelihood estimates of coefficients for the distribution, accomodating for variability across samples. Finally, DEG testing was performed using a likelihood ratio test in the ‘glmLRT’ function with a contrast matrix defined by the ‘makeContrasts’ function to compare between smoker and non-smokers. Analysis results were FDR-corrected by Benjamini-Hochberg method using the ‘topTags’ function. These results were compiled for each cell phenotype and the final output exported to a .csv file. To complement the prior analysis, cell phenotypes most perturbed to smoking status were investigated using the Augur package (v1.0.3). Scaled and normalized RNA counts were used to calculate areas under the curve per phenotype with the ‘calculate_auc’ function, with higher numbers representing higher biological perturbations within a high-dimensional space quantified using a machine learning classifier framework.

##### Trophoblast trajectory modelling

To model the differentiation trajectory of the trophoblast, subsetting of the 67,618 nuclei annotated as trophoblast cell types or states (CTBp, CTB, CTBpf, STBim, STB, CCT) was performed. To ensure a most adequate inference, gene expression counts were further filtered to include nuclei with a minimum count of 600 genes, resulting in 63,077 nuclei included for this analysis. First, the non-integrated gene expression count matrix was split into a test and training matrices of counts using the ‘countsplit’ function of the countsplit package (epsilon 0.5, v1.0.0). These matrices are independent under specific modelling assumptions and are therefore robust for use in cross validations. The trajectory modelling was performed on the test matrix and the leaf and transition gene analysis on the training matrix. Linear dimensionality reduction was performed on the 4,000 most variable genes of the integrated dataset and 30 PCs were calculated. The K-nearest neighbour graph was computed using the first 30 PCs. UMAP visualisation was used to further reduce the high dimensional latent spaces to 2D using the first 30 PCs. Louvain clusters were computed in an unsupervised manner based on the k-nearest neighbour graph at resolutions between 0.1 and 1 in 0.1 step increments using the Louvain algorithm with modularity optimizer version 1.3.0 by Ludo Waltman and Nees Jan van Eck, including a random seed for reproducibility. The most appropriate resolution to use for annotation was selected based on cluster relationship visualisation at all resolutions by building a cluster tree using the clustree package (v0.5.1). The resolution of 0.3 yielding 10 clusters was used. Counts and associated metadata from Seurat were used to create a SingleCellExperiment object (v.1.22.0). To evaluate if a shared or separate trajectory should be fit depending on maternal smoking staus, an imbalance score was calculated using the ‘imbalance_score’ function (k = 20, smooth = 40) from the condiments package (v1.2.0). Given that no score was greater than 3.5, a common trajectory was fitted. The trajectory was modelled on the subsetted UMAP graph using the ‘slinghot’ function in the slingshot package (v2.2.1) with the starting cluster set to cluster 2 based on its expression of proliferating markers *TOP2A* and *MKI67*. The function identifies global structure with a cluster-based minimum spanning tree and fits simultaneous principal curves to describe each lineage. A pseudotime is allocated to each nucleus for a maximum of two possible trajectories, with an associated weight indicating the predicted certainty of each assignment. Modelled trajectories included three distinct lineages going towards the STB fate, the CTB fate and the CCT fate. Differences between pseudotime densities for each lineage were assessed in consideration of the curve weights, by a permutation test (10,000 replicates) and Kolmogorow-Smirnov per lineage using the ‘progressionTest’ function of the slingshot package. Differential differentiation between smoking status along pseudotime per lineage was assessed by a quasibinomial generalised linear model using the ‘glm’ function in the stats package (v4.1.2).

Transition and leaf gene inference was performed on non-smokers nuclei only (*n* = 36,022 nuclei). Transition genes along each inferred lineage trajectory were calculated as described in Chen et.al.^86^. First, genes with a log_2_FC lower than ± 0.25 between the first 20% and last 20% of nuclei in the inferred pseudotime were filtered out. For transition genes, expression values per modelled lineage were scaled between 0 and 1, and a spearman correlation between gene expression and pseudotime values was fitted gene-wise using the ‘cor.test’ function of the stats package (v4.1.2). Genes with a Spearman’s correlation coefficient of ± 0.4 were identified and reported as transition genes. For the leaf gene calculation, DEGs per lineage were empirically calculated using the edgeR package (v3.36.0). A DGEList object from the count matrix was created and genes that were not expressed above a log-CPM threshold in a minimum of five samples were filtered out. Normalization factors were calculated using the trimmed mean of M-values (TMM) method to account for composition biases in the data. A design matrix was constructed and included batch and fetal sex to model the effects of these variables on gene expression. Robust dispersion estimates were obtained using the ‘estimateGLMRobustDisp’ function, which accommodates for potential outliers. A negative binomial generalized linear model was fitted genewise using the ‘glmFit’ function with robust parameter settings to compute the maximum likelihood estimates of coefficients for the distribution, accomodating for variability across samples^151^. Finally, DEG testing was performed using a likelihood ratio test with the ‘glmLRT’ function with a contrast matrix defined by the ‘makeContrasts’ function to compare between smoker and non-smokers. The results of the analysis were FDR-corrected by Benjamini-Hochberg method using the ‘topTags’ function.

##### Network hub genes inference, TF-prediction and pathway enrichment analyses

The list of DEGs based on cut-off values (logFC +/-0.25 and FDR < 0.1) were used as background for networks. Genes were used as input in the stringDB for protein-protein interaction (PPI) networks (confidence level = 0.4, no added proteins in shells). Networks were then further analysed in Cytoscape (v3.10.1). Hub genes, defined as genes with high connectivity across DEGs, were identified from the PPI network calculating top 10 genes for all CytoHubba plug-in topological analysis methods (DMNC; MNC, MCC, ecCentricity, Bottleneck, Degree, EPC and Closeness). The candidate hub genes were merged into one network and decomposed into communities using the cytoscape plug-ins clustermaker and GLay, based on Newman and Girvan’s edge-betweenness algorithm. The original background log_2_FC was used for continuous mapping colours. The hub gene network was used to calculate transcription factors via the plug-in iRegulon (standard threshold: enrichment score threshold 3.0, ROC threshold for AUC calc 0.03, Rank threshold 5000, minimum identity between orthologous genes: 0.0, max FDR on motif similarity: 0.001). Predicted transcription factors were visualised as PPI (confidence level 0.15, no added proteins in shells) via stringDB.

Pathway enrichment analysis was performed using the 1D annotation enrichment method in Perseus software (v1.5.15.0). Enrichment was done based on ranked log fold changes without thresholding spanning the gene set enrichment analysis (GSEA), reactome and the WikiPathways databases.

#### 3.3.2. Proteomics

Proteomics measurements were analysed using the timsControl software (Bruker Daltonik GmbH, v. 3.1). Raw files were analysed with DIA-NN (v. 1.8.1 and 1.8.2 beta 25; for DVP and cell pellet samples, respectively)^163^ in library-free mode based on a predicted human spectral library (Uniprot 2021 release). Default settings were used with small adjustments. The mass range was set to 100 – 1,700 m/z, precursor charge state was 2 - 4 and the maximum number of allowed miscleavages was 2. MS1 and MS2 mass accuracies were set to 15 ppm and the match-between-runs option was enabled. Quantification strategy was set to ‘Robust LC’. For downstream data analysis, we used the protein FDR filtered pg_matrix.tsv and unique_genes_matrix.tsv DIA-NN output tables were analysed with Perseus (v. 1.6.15.0) and the Protigy interactive web app (v. 1.0.2, https://github.com/broadinstitute/protigy). Missing values were imputed based on a shifted normal distribution of log_2_ transformed sample values (width = 0.3; downshift = -1.8) after stringent data filtering (minimum of 70% quantified values across samples). In the case of microdissected cells for DVP, prior to principal component analysis (PCA), batch effects were corrected with the proBatch R package (v. 1.10.0) based on the ComBat method. Differential abundance was evaluated using the limma package (v 3.56.2) by fitting a linear model (lmFit) on a design matrix of protein abundance data adjusted by a contrast matrix representing smoking status. Empirical Bayes moderation (eBayes) was used on the standard errors to produce moderated t-statistics. False discovery rate was controlled using a Benjamini-Hochberg correction. Pathway enrichment analysis was performed using the 1D annotation enrichment method in Perseus software. Enrichment was done based on ranked log fold changes spanning the gene ontology (GO), gene set enrichment analysis (GSEA), reactome and the Kyoto encyclopedia of genes and genomes (KEGG) databases.

##### HubGene inference and transcription factor prediction

The list of differentially abundant proteins based on cut-off values (logFC +/-0.25 and FDR < 0.1) were used as background for networks. Proteins were used as input in the stringDB for protein-protein interaction (PPI) networks (confidence level = 0.4, no added proteins in shells). Networks were then further analysed in Cytoscape (v3.10.1). Hub genes, defined as genes with high connectivity across differentially abundant proteins, were identified from the PPI network calculating top 10 genes for all CytoHubba plug-in topological analysis methods (DMNC; MNC, MCC, ecCentricity, Bottleneck, Degree, EPC and Closeness). The candidate hub genes were merged into one network and decomposed into communities using the cytoscape plug-ins clustermaker and GLay, based on Newman and Girvan’s edge-betweenness algorithm. The original background log_2_FC was used for continuous mapping colours. The hub gene network was used to calculate transcription factors via the plug-in iRegulon (enrichment score threshold 3.0, ROC threshold for AUC calc 0.03, Rank threshold 5000, minimum identity between orthologous genes: 0.0, max FDR on motif similarity: 0.001). Predicted transcription factors were visualised as PPI (confidence level 0.15, no added proteins in shells) via stringDB.

Pathway enrichment analysis was performed using the 1D annotation enrichment method in Perseus software (v1.5.15.0). Enrichment was done based on ranked log fold changes without thresholding spanning the gene set enrichment analysis (GSEA), reactome and the WikiPathways databases.

For in-vitro and validations, groups were compared between smoking and non-smoking groups using two-tailed Mann-Whitney U test in GraphPad Prism software (v10.2.0).

#### 3.3.3. Visualizations

The following packages were used to compile the computational analyses visualisations: tidyr (v1.2.0), ggplot2 (v3.3.6), patchwork (v1.1.1), cowplot (v1.1.1), enhancedVolcano (v1.18.0), viridis (v1.6.4), ComplexHeatmap (v2.16.0), RColorBrewer (1.1.3), Seurat (v4.1.0), RVenn (1.1.0), slingshot (v.2.2.1), clustree (v.0.5.1), pheatmap (v1.0.12). Baseline characteristics, qPCR and ELISA results were plotted using GraphPad Prism (v10.2.0). Results from network analyses (hub gene and pTF inference) were visualised using cytoscape (v3.10.1).

Figure panels were arranged using Adobe Illustrator (v27.5).

